# Shifting KRAS hotspot mutations inhibition paradigm in colorectal cancer

**DOI:** 10.1101/2023.08.09.552513

**Authors:** Ana Rita Brás, Ana Lopes, Nuno Mendes, Paulo J. Costa, Anabela Ferreira, Sara Granja, Ana Paula Silva, Francisco Tortosa, Fátima Baltazar, Fátima Gärtner, Maria João Sousa, Andreia Valente, Ana Preto

**Affiliations:** Centre of Molecular and Environmental Biology (CBMA), Department of Biology, University of Minho, Campus de Gualtar, 4710-057, Braga, Portugal; Institute of Science and Innovation for Bio-Sustainability, University of Minho, Campus de Gualtar, 4710-057, Braga, Portugal; Centro de Química Estrutural, Institute of Molecular Sciences and Departamento de Química e Bioquímica, Faculdade de Ciências, Universidade de Lisboa, Campo Grande, 1749-016 Lisboa, Portugal; Institute for Research and Innovation in Health (i3S), University of Porto, 4200-135 Porto, Portugal; Institute of Molecular Pathology and Immunology of the University of Porto, 4200-135 Porto, Portugal; BioISI - Instituto de Biosistemas e Ciências Integrativas and Departamento de Química e Bioquímica, Faculdade de Ciências, Universidade de Lisboa, 1749-016, Lisboa, Portugal; Department of Pathological, Cytological and Thanatological Anatomy, School of Health, Polytechnic Institute of Porto, 4200-072 Porto, Portugal; Life and Health Sciences Research Institute (ICVS), School of Medicine, University of Minho, 4710-057 Braga, Portugal; ICVS/3B’s-PT Government Associate Laboratory, 4710-057 Braga, Portugal; ICVS/3B’s-PT Government Associate Laboratory, 4806-909 Guimarães, Portugal; Center for Interdisciplinary Research Egas Moniz (CiiEM), Quinta da Granja, Monte da Caparica, 2829-511 Caparica, Portugal; Instituto de Anatomia Patológica, Faculdade de Medicina da Universidade de Lisboa, Av. Prof. Egas Moniz, 1649-028 Lisboa, Portugal; Institute of Biomedical Sciences of Abel Salazar, University of Porto, 4050-313 Porto, Portugal

**Author notes:** These authors share senior authorship (Andreia Valente, Ana Preto).

**Keywords:** PMC79, colorectal cancer, KRAS inhibition, KRAS^G12V^ mutations, KRAS^G12D^ mutation, KRAS^G13D^ mutation

## Abstract

KRAS hotspot mutations are difficult to target, highlighting the need of developing new specific target drugs for cancers driven by these mutations, like colorectal cancer (CRC). Here, we discover a new ruthenium compound, PMC79, that inhibits specifically mutated KRAS and the downstream signaling ERK and AKT proteins both “in vitro” and “in vivo”. We demonstrated that PMC79 inhibits KRAS mutated kinase activity and is selective for KRAS mutations not affecting the KRAS wild-type protein. KRAS inhibition is not dependent on actin polymerization or on proteasome. Molecular docking analysis suggests that this effect might result from protein dynamics associated with the mutations. We demonstrated that low doses of PMC79 potentiate 5-fluorouracil anticancer effect. “In vivo” PMC79 “proof of concept” showed that it reduces tumor growth in the CAM-xenograft model and induces necrosis of the tumor in the xenograft mice model. PMC79 is a promising new “magic bullet” for CRCs harboring mutated KRAS.

## 1. Introduction

Targeting KRAS mutations has been a clinical challenge all over the years due to its great relevance and role in cancer, being for many years called the “undruggable KRAS”. Activating mutations in KRAS oncoprotein are a hallmark of cancer being a key player in intracellular signaling pathways regulating the carcinogenesis process^1^. KRAS mutations are present in around 21.6% of human cancers^2,3^, being most frequently observed in pancreatic ductal adenocarcinoma (PDAC) (80%), colorectal cancer (CRC) (40%) and non-small cell lung cancer (NSCLC) (35%)^4^ and also in 10% of ovarian and endometrial cancers^5^.

KRAS hotspot mutations are frequently associated with alterations at codons 12 and 13^6^, which result in G12D, G12V or G13D mutations. A special case occurs in NSCLC where a switch from glycine to cysteine results in G12C mutation^7^. This hotspot mutation account for approximately 40% of the 35% of the NSCLC cases with KRAS mutation^8^. All these alterations lead to conformational changes in KRAS protein structure resulting in its constitutive activation and the activation of the downstream MAPK and PI3K signaling pathways^9,10^.

CRC is among the most prevalent type of cancer being the second cause of cancer death around the world^11^. CRC is the second cancer type with the highest presence of KRAS hotspot mutations being G12D (13%), G12V (9%), and G13D (7%) the most frequently found^4^. There are limited therapeutic approaches for CRC bearing KRAS mutations as they are resistant to all the available therapies including anti-EGFR target therapy drugs such as EGFR antibodies bevacizumab, cetuximab^12,13^. The resistance of CRC harboring KRAS mutations to these inhibitors creates a clinically relevant problem that needs to be overcome.

For many years, KRAS was considered an “undruggable” target, however, in recent years this perspective has changed with the appearance of several molecules capable of inhibiting this protein^14^.

Despite all the efforts performed over the years, specific targeting of KRAS hotspot mutations G12V, G12D and G13D has been very difficult to achieve. The lack of specific anticancer agents for targeting KRAS in CRC bearing this mutation highlights the need of developing new KRAS hotspot mutations specific target drugs. In the last years, two KRAS^G12C^ inhibitors, sotorasib and adagrasib, have been approved by the FDA for the treatment of NSCLC with KRAS^G12C^ mutations^14^. These inhibitors selectively form a covalent bond with cysteine 12 within the switch-II pocket of KRAS^G12C^ protein, thereby locking KRAS in the inactive state^15–21^. However, KRAS^G12C^ mutation is only present in 3% of CRCs which makes these inhibitors ineffective against CRCs with mutated KRAS^22^. Recent studies have identified a potent and selective KRAS^G12D^ inhibitor, MRTX113, whose antitumor benefit promoting tumor regression was only demonstrated in a murine autochthonous pancreatic cancer model^23,24^. Despite some KRAS inhibitors have entered clinical trials, most of them target KRAS^G12C^ mutations^21,25–28^ and these were the only ones approved for clinical use. To the best of our knowledge, no KRAS inhibitors are available in the clinics to specifically target all KRAS hotspot mutations present in CRC.

Metal complexes in cancer therapy have attracted high interest over the years^29–31^. Since the successful discovery of cisplatin in 1965, a new era in the metal-based anticancer drugs history has emerged^32^. Cisplatin and its derivatives, oxaliplatin and carboplatin, are the most studied group of metallodrugs and have been broadly used to treat several cancer types, including CRC^33^. However, the numerous side effects and drug resistance problems, and the absence of specific targets have limited their use^34^. Recently, several other metal complexes emerged as alternatives to platinum drugs, namely based on zinc, copper, iron, ruthenium, gold, and silver, among others^32^. Some of these metal-based compounds are already in clinical trials and several more are awaiting ethical approval to enter the trial. Metal-based compounds are recognized to have a high therapeutic potential mainly due to their redox characteristics, variable coordination modes, and reactivity with organic substrates^35^. These properties increase the interest in the design of metal complexes that specifically target biomolecules in cancer cells. The increased specificity and selectivity for cellular targets as membrane receptors and transporters, intracellular enzymes, nucleic acids and organelles has gained a lot of attention from researchers and has been an area of high progress in recent years^36–38^.

Of the several known metallodrugs, ruthenium compounds are one of the most investigated concerning their anticancer effects and mechanisms of action, as well as clinical evaluation^39–43^. Currently, there are two complexes in clinical trials: NKP1339 for CRC treatment and photodynamic therapy with TLD1433 for nonmuscle invasive bladder cancer treatment^39,44,45^.

Over the last years, our group has been studying a family of “piano stool” compounds, named Ru(II)-cyclopentadienyl complexes, as potential anticancer drugs for chemotherapy^43^. The general structure of these complexes is based on [Ru(η^5^-C_5_H_5_)(PPh_3_)(2,2’-bipy)]^+^ structure and its derivatization. Several compounds of this family showed good anticancer characteristics against a wide panel of cancer cell lines^46–58^. “In vivo” studies in embryo larval zebrafish model showed that some compounds are well tolerable showing promising anticancer properties^50,58^. The mice model results showed some compounds were tolerant to the maximum dose tested, despite some liver toxicity^49^. Moreover, one of the compounds, LCR134, proved to be a specific P-gp inhibitor^50^.

In this work, we discover for the first time that the organometallic Ru compound, PMC79 [Ru(η^5^-C_5_H_5_)(PPh_3_)(2,2’-bipy-4,4’-CH_2_OH)]^+^ (Figure 1a), has a specific effect in inhibit KRAS mutations in several CRC models with different KRAS hotspot mutations. Previous “in vitro” studies showed that PMC79 is highly active in CRC cell lines, inhibiting proliferation, migration and inducing apoptosis^46,47^. Furthermore, “in vivo” studies demonstrated that this compound has an anti-proliferative capacity in the zebrafish model^50,58^.

**Figure 1.**
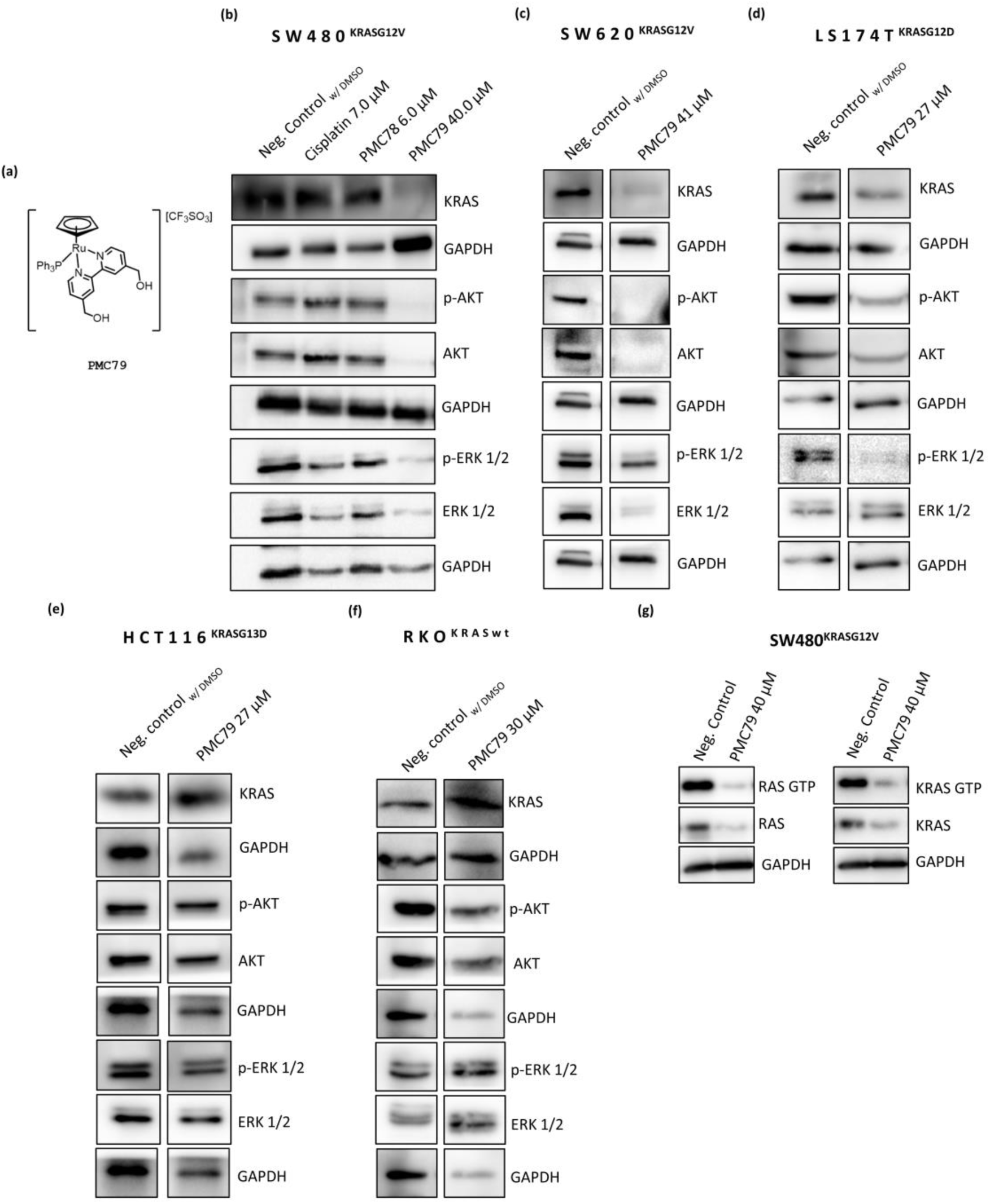
PMC79 decreases the expression of KRAS and KRAS downstream signaling molecules in CRC cells with KRAS mutation. **(a)** PMC79 chemical structure. **(b)** Immunoblot analysis of KRAS signaling molecules in SW480 cells after 48 h treatment with cisplatin, PMC78 and PMC79. **(c-f)** Immunoblot analysis of KRAS signaling molecules in SW620, LS174T, HCT116 and RKO cell lines, respectively, after 48 h treatment with PMC79. The results were obtained from at least three independent experiments. **(g)** Immunoblot analysis of RAS-GTP and KRAS-GTP in SW480 cells after 48 h treatment with PMC79. The results were obtained from two independent experiments.

We aimed to go further on the quest of identifying the mechanism of action of PMC79 analyzing its effect on KRAS signaling pathways in CRC harboring KRAS mutations. To our surprise, to the best of our knowledge we discover for the first time a Ru organometallic compound that specifically targets KRAS hotspot mutations G12V, G12D, G13D and MAPK and PI3K signaling pathways. PMC79 provides a more direct approach to selectively inhibit oncogenic KRAS mutant function while sparing the function of KRAS wild-type protein in CRC. The new Ru compound PMC79 is the first specific inhibitor of KRAS hotspot mutations identified, constituting a promising drug for the treatment of resistant CRC with KRAS mutations.

## 2. Experimental section

### 2.1 “In vitro” studies

#### 2.1.1 Compounds and reagents

PMC79, PMC78, LCR134 and LCR220 were synthesized as previously reported by us^49,50,52^. Cisplatin, latrunculin A, MG132 (M7449, Sigma-Aldrich, St. Louis, MO, USA), adagrasib (T8369, TargetMol, Wellesley, MA, USA), sotorasib (T8684, TargetMol, Wellesley, MA, USA) and 5-Fluorouracil (5-FU) (F6627, Sigma-Aldrich, St. Louis, MO, USA) were dissolved in dimethyl sulfoxide. Cisplatin, MG132, adagrasib and sotorasib were stored at −20 °C, and latrunculin A and 5-FU were stored at −80 °C and 4°C, respectively.

#### 2.1.2 Cell Lines and Culture Conditions

CRC-derived cell lines SW480^KRASG12V^, SW620^KRASG12V^, LS174T^KRASG12D^, HCT116^KRASG13D^ and RKO^BRAFV600E; KRASwt^, were obtained from American Type Culture Collection (Manassas, VI, USA). SW480 and SW620 cell lines grew in Roswell Park Memorial Institute 1640 medium with stable glutamine (Biowest, Nuaillé, France); LS174T and RKO cell lines grew in Dulbeccós Modified Eaglés Medium High Glucose (Biowest, Nuaillé, France); HCT116 cell line grew in McCoy’s 5A Medium (Biowest, Nuaillé, France). All mediums were supplemented with 10% fetal bovine serum (v/v) (Biochrom, Berlin, Germany) and 1% penicillin/streptomycin (v/v) (Biowest, Nuaillé, France). All cell lines were maintained at 37 °C under a humidified atmosphere containing 5% CO_2_.

#### 2.1.3 Cell Treatments

SW480, SW620, LS174T, HCT116 and RKO cell lines were seeded at 1×10^5^ cells/ml, 1.3×10^5^ cells/ml, 7.5×10^4^ cells/ml, 1×10^5^ cells/ml and 1×10^5^ cells/ml respectively and adhered onto appropriate sterile plates. In some specific assays, SW480 cells were seeded at 7.5×10^4^ cells/ml and 6×10^5^ cells/ml. 24 h after seeding, cells were incubated with the corresponding treatments in all experiments. For the experiments using the proteasome inhibitor MG132, cells were pretreated with 10 μM of MG132 for 1 h and then co-treated with PMC79 for 8 h and 12 h.

#### 2.1.4 Sulforhodamine B Assay

Sulforhodamine B (SRB) assay was performed to determine the half-maximal inhibitory concentration (IC_50_) of PMC79 in SW620, LS174T and HCT116 cell lines and adagrasib, sotorasib and 5-FU in SW480 cell line, after 48 h of treatment.

Cell lines were seeded in 24-well plates and after 24 h were incubated with different concentrations of the compounds for 48 h. SRB assay was performed as previously described^46^. Results were expressed relative to the negative control, which was considered as 100% of cell growth. The IC_50_ was estimated using GraphPad Prism 8 software, applying a sigmoidal dose vs response (variable slope) non-linear regression (n=3).

#### 2.1.5 Active Ras pull-down assay

SW480 cells were seeded in 100 mm Petri dishes and treated with different concentrations of PMC79. After 48 h, RAS activity was determined by the Active Ras Pull-Down and Detection Kit according to the manufacturer’s instructions (16117, Thermo Fisher Scientific, Lafayette, CO, USA).

#### 2.1.6 KRAS RNA silencing

Small interference RNA (siRNA) targeting human KRAS was designed by Horizon Discovery (L-005069-00-0020, Horizon Discovery, Cambridge, UK). SW480 cells were seeded in 6-well plates and after 24 h, cells were transfected with 7.5 nM siRNA against KRAS. Transfection quality was monitored using a validated control (non-silencing) siRNA (1027281, Qiagen, Germantown, MD, USA). SW480 cells were transfected according to the manufacturer’s instructions, using 3 μL of lipofectamine 2000 (11668019, Invitrogen, Waltham, MA, USA). Control cells (blank) were left untreated. After 6 h, the transfection mixture was removed, and cells were left untreated or treated with PMC79 and incubated for a further 48 h in fresh medium. After 48 h, samples were processed as described in section 2.1.9 for immunoblot analysis.

#### 2.1.7 Actin polymerization inhibition using latrunculin A

Primarily, the effect of latrunculin A on cell viability was determined by the SRB assay in the SW480 cell line according to the procedure described in section 2.1.4. In addition, the alterations of F-actin were assessed using Alexa Fluor^TM^ 568 Phalloidin by fluorescence microscopy as reported by us ^46^. After the determination of the latrunculin A dose that inhibits actin polymerization without causing a decrease of cell growth, SW480 were co-incubated with 75 nM of latrunculin A and 40 μM PMC79 for 48 h. Then, cells were fixed and stained using Alexa Fluor^TM^ 568 Phalloidin. Furthermore, cells were also collected for protein extraction and immunoblot analysis as described in section 2.1.9.

#### 2.1.8 KRAS-humanized Saccharomyces cerevisiae model

*Saccharomyces cerevisiae (S. cerevisiae)* W303-1A strain, deleted in *RAS2*, the most expressed yeast RAS isoform, was transformed with human KRAS isoforms, KRAS^wt^, and the most common KRAS mutations in CRC, KRAS^G12D^, KRAS^G12V^ and KRAS^G13D^, using the pCM184 plasmid. pCM184 yeast plasmid (*TRP1*) is under the control of a tet-off promoter, indicating that the addition of doxycycline leads to its repression^59^.

##### 2.1.8.1 Saccharomyces cerevisiae growth conditions

*S. cerevisiae* cells were grown under aerobic conditions in synthetic complete medium (SC: 0.17% (w/v) yeast nitrogen base without amino acids, 0.5% (w/v) ammonium sulfate, 2% (w/v) glucose, 0.2% (w/v) dropout mixture without the auxotrophic marker tryptophan). Solid medium was obtained through the addition of 2% (w/v) agar. All incubations were performed under the same conditions: 30 °C and 200 r.p.m.

##### 2.1.8.2. Saccharomyces cerevisiae viability assay

Cells were grown in SC medium, at 30 °C and 200 rpm, until exponential phase. Then, cells were resuspended in fresh medium to a final OD_640nm_ of 0.3, and PMC79, PMC78 and Adagrasib were added at different concentrations. DMSO was used as a negative control (maximum of 0.3% (v/v)). Cell viability was assessed at time 0 and after 1, 2, 4, 8 and 24 h. Cells were plated into YPD agar plates (YPD: 1% (w/v) yeast extract, 2% (w/v) peptone, 2% (w/v) glucose, 2% (w/v) agar) after serial dilutions of 1:10. After incubation for 2 days at 30 °C, the number of colony forming units (CFUs) was counted and the percentage of viable cells was calculated, considering the number of CFUs at time 0, right before the addition of compounds, as 100%.

#### 2.1.9 Immunoblot analysis

Preparation of total protein extracts of human cell lines was performed as previously described in ^55^. Preparation of total protein extracts of yeast cells was initiated after harvesting cells and resuspending in 500 µL of deionized water. First, cells lyse was performed through the addition of 50 µL of 3.5% (v/v) β-mercaptoethanol in 2 M NaOH solution, and incubation for 15 minutes, in ice. Then, 50 µL of 50% trichloroacetic acid (TCA) was added in order to precipitate proteins. After incubation in ice for 40 minutes to 1 hour, cells were centrifuged at 12000 g for 10 minutes. Samples were washed with acetone and then solubilized in a solution containing 1 M NaOH and 2% (v/v) SDS. Finally, the Laemmli buffer (4x: 0.25 M Tris-HCl, 9.2% (w/v) SDS, 40% (w/v) glycerol, 5% (w/v) β-mercaptoethanol, 0.5% (w/v) bromophenol blue) was added and samples were denatured at 70 °C for 15 minutes. SDS-PAGE and Western blots were performed as described in ^55^.

#### 2.1.10 Antibodies

Antibodies used were: anti-KRAS and anti-β-actin (Sigma, St. Louis, MO, USA); anti-phospho p44/42 MAPK (Thr202/Tyr204) (pERK), anti-p44/42 total (ERK), anti-phospho Akt (Ser473) (pAKT) and anti-Akt total (Cell Signaling, Danvers, MA, USA) (AKT); anti-RAS (Thermo Fisher Scientific, Lafayette, CO, USA); GAPDH (Gene Tex, Irvine, CA, USA); anti-yeast phosphoglycerate kinase (PGK1) (Molecular Probes, San Jose, CA, USA).

#### 2.1.11 Statistical analysis

The results were obtained from at least three independent experiments and expressed as mean ± SD. A one-way ANOVA with Šídák’s multiple comparisons test and a two-way ANOVA with Dunnett’s post-test were used to analyze the results. *p*-values lower than 0.05 were considered statistically significant. All statistical analyses were performed using GraphPad Prism version 8.

### 2.2 “In silico” studies

#### 2.2.1. System construction and molecular dynamics simulations

Herein we followed an ensemble docking approach^60,61^ in which the conformation selection is made by the ligand among an ensemble of conformations of the apo target. To generate such ensemble we used molecular dynamics (MD) simulations of the target protein. From the Protein Data Bank (PDB) we selected 3 crystal structures of the GDP(Mg^2+^)-bound KRAS featuring the two mutations of interest, G12D (PDB: 4EPR)^62^ and G12V (PDB: 7C40)^63^, along with the wild-type (PDB: 4OBE)^64^. Protonation states at pH 7.0 were assigned using PypKa^65^ and then, MD simulations were carried out using the GROMACS software package, version 2020.6^66^ along with the AMBER99SB-ILDN force field^67^ for the protein while parameters for GDP were taken from reference^68^. The guanosine diphosphate (GDP), the magnesium cation (Mg^2+^) and the crystallographic water molecules were kept in their X-ray positions in the starting structure. A dodecahedral simulation box was used with a distance between the solute and the box of 1.0 nm while applying periodic boundary conditions in all directions with the minimum image convention. The systems were solvated with TIP3P water molecules and neutralized with sodium cations. The electrostatic interactions were treated with particle mesh Ewald (PME)^69^. Here, a short-range electrostatic cutoff of 1.0 nm was used along with a Fourier grid spacing of 0.125 nm. A 1.0 nm cutoff was used for the van der Waals interactions whereas bonds to the hydrogen atoms were constrained using the P-LINCS algorithm^70^. The systems were initialized by performing a two-step energy minimization using the steepest descent algorithm, the first without constraint, and the second using constraints on bonds involving hydrogen atoms. Then, an NVT simulation was run for 100 ns keeping the temperature at 298.15 K using the Berendsen thermostat^71^ followed by 100 ps of NPT simulation using the v-rescale thermostat^72^ while the Parrinello– Rahman barostat^73^ was used to maintain pressure at 1 bar. Then, a production run was performed during 200 ns. The first 20 ns of this NPT run were discarded as equilibration and then, 180 equally-spaced snapshots were extracted for each system.

#### 2.2.2 Molecular Docking

The ensemble of receptor structures, taken from the MD simulations, were RMSD fitted and converted to the PDBQT format using the *prepare ligand* tool of ADFRsuite version 1.1^74^. In this step, the non-polar hydrogen atoms were removed and AutoDock atom types were assigned. The GDP and magnesium were kept in the receptor structure. A tridimensional model of PMC79 was built and geometry optimized with the PBE1PBE functional^75^ using Gaussian09. The 6-31G* basis set was used for all elements apart from ruthenium and phosphorus for which the LANL2TZ(f) basis set along with an effective core potential was employed. The optimized structure was converted to the PDBQT format using AutoDockTools, also removing non-polar hydrogen atoms, assigning AutoDock atom types and the allowed torsions. Parameters for ruthenium are not available, therefore, following previous reports^76,77^, this element was replaced by iron while keeping the ruthenium-optimized distances. Such substitution is not problematic since the metal is not directly exposed to the protein.

The Molecular Docking simulations were performed with AutoDockVina 1.2.5^78^. For each system (G12D, G12V, WT) and receptor conformation, compound PMC79 was docked using Vina and Vinardo^79^ scoring functions. A search space of 25 × 25 × 25 Å was used, centered near Switch-II (see Figure S1, green dot). This search space is wide enough as to allow the ligand to explore not only Switch-II (Figure S1, blue) but also the regions near the phosphate-binding loop (P-loop, yellow in Figure S1) and Switch-I (green in Figure S1). Notice the P-loop is responsible for GDP binding whereas Switch-I and Switch-II are key regions for the binding of small molecules. All protein-ligand poses were then sorted according to the Vina and Vinardo scoring functions, thus allowing to select the best conformation to bind PMC79.

### 2.3 “In vivo” studies

#### 2.3.1 “In vivo” chick embryo chorioallantoic membrane assay

Chick embryo chorioallantoic membrane (CAM) assay was performed as described in ^80^. On the 9^th^ day of development, SW480 cells (2×10^6^ cells) were mixed with matrigel and placed inside the CAM. After 4 days (day 13), tumors were treated with PMC79 (25.0 μM and 40.0 μM) and 5-FU (12.6 μM and 73.5 μM). “In ovo*”* tumor area analysis was performed using ImageJ software. “Ex ovo” images were used to obtain blood vessel analysis using Fiji software with “Vessel analysis” plugin.

##### 2.3.1.1 Statistical analysis

The results were obtained from at least 10 eggs per experimental condition and expressed as mean ± SEM. A one-way ANOVA with Dunnett’s multiple comparisons test was used to analyze the results. *p*-values lower than 0.05 were considered statistically significant. All statistical analyses were performed using GraphPad Prism version 8.

#### 2.3.2 “In vivo” studies in nude mice

Animal experiments were previously approved by the Animal Ethics Committee and the animal welfare body of Institute for Research and Innovation in Health (i3S) and the Directorate General of Food and Veterinary and were carried out in accordance with all the guidelines of the Portuguese Society for Science in Laboratory Animals and the European Guidelines for the Care and Use of Laboratory Animals, Directive 2010/63/EU.

In all “in vivo” experiments female N:NIH(S)II-nu/nu mice, strain described by Azar et al. in 1980^81^, were used. The mice were bred, housed and maintained at i3S Animal House in a pathogen-free environment under controlled conditions of light, temperature and humidity. The following Humane Endpoints for euthanasia were established: i) any signals of distress, suffering or pain; ii) weight loss greater than 20-25% of the body mass; and iii) anorexia and moribund state, related or not to the experimental procedure. In all experiments, mice aged 6-9 weeks old were used and were monitored at least 4 times per week.

##### 2.3.2.1 “In vivo” toxicity in a colorectal cancer mice xenograft model

To evaluate both the lethal and the tolerated doses of PMC79 in a formulation with 40% Captisol® in Milli-Q water (m/v), five doses were tested: 34, 26, 17, 13 and 8.7 mg/Kg per mice (n = 4 per group, except the last one in which n = 2). Mice were intraperitoneally injected with PMC79 three times a week for 17 days. The control group received injections of 450 µl of vehicle. Animal reaction to PMC79 was observed immediately after the injection and mice were monitored and weighed every day throughout the study period. On day 19, the animals were euthanized through cervical dislocation after being anesthetized by intraperitoneal (IP) injection of a solution of 150 mg/Kg of ketamine with 2 mg/Kg of medetomidine.

##### 2.3.2.2 Biochemical blood analysis

On day 19, the animals from the toxicological study were anesthetized for whole venous blood collection by intracardiac puncture. After collection, the blood was placed in microtubes with about 20 µL of heparin each. The microtubes were then placed in a centrifuge for 20 minutes at 3500 rpm, after which the precipitate was discarded, and the supernatant (plasma) was collected and placed in a new microtube. The plasma samples were further processed for biochemical analysis.

##### 2.3.2.3 “In vivo” anti-tumor activity in a xenografted colorectal cancer mice model

A total of 16 female N:NIH(S)II-nu/nu mice were heterotopically inoculated with 1×10^6^ viable SW480 cells in the right side of the animal’s back, slightly lateral to the scapula using a 25 g needle. As soon as the nodules were visible (day 3), mice were randomized and divided into 2 groups: PMC79 treatment group (n=8; Group 1) and control (n=7; Group 2). Group 1 received IP injections of 17 mg/Kg of PMC79 while Group 2 was injected with the vehicle. Mice were treated accordingly to Table S1, and tumors were surgically removed on days 20-32 post-inoculation as they reached an average volume of approximately 1000 mm^3^. Tumor size was measured using calipers, and tumor volumes (mm^3^) were estimated using the formula: W x L^2^ x ½, where W is the width and L is the length of the tumor. For tumors removal, mice were anesthetized using an anesthesia machine (VetTech Solutions Ltd): anesthesia was first induced in an induction chamber with 5% isoflurane and 1-2% oxygen and then anesthesia was maintained by decreasing isoflurane to 1.5-2%. After anesthesia, a total excision of the tumors was performed. To avoid pain, the mice were subcutaneously treated with a solution of 0.08 mg/Kg of buprenorphine per mouse (100 μL/10 g of animal) before and after tumor excision and afterward twice a day for 48 h. At the time of surgery, a small piece of the tumor was placed in liquid nitrogen for protein extraction and analysis of the KRAS expression. The remaining tumor was fixed in 10% buffered formalin for histology. The animals that underwent surgery were not given further treatment, and 5-7 days after tumor removal, a weekly maintenance dose of the respective treatment began to be administered. All animals were euthanized on day 67 post-inoculation (35 days after the removal of the last tumors), being anesthetized using a tube with isoflurane. Lungs and lymph nodes (4 submandibular, 4 axillary and 2 inguinal) were collected from each animal and immediately fixed 10% buffered formalin for histology to search for micrometastases.

##### 2.3.2.4. Histopathological and immunohistochemical analysis

The tissues were fixed in buffered formalin and the inclusion in paraffin was done according to the usual technical procedures. Histological sections of 3 microns, stained with hematoxylin and eosin (H&E) and mounted on microscope slides were made. The slides were observed with an optical microscope (Zeiss Axioskop 2) and images were captured using a digital camera coupled (DS Camera Control Unit DS-L2). Histological evaluation was made in blind analysis by two pathologists. The samples were compared with those of control mice and histological variables as necrosis, intratumoral hemorrhage, lymphocytic infiltration, peritumoral oedema, as well as the degree of tumor regression were evaluated. The proliferative index of the tumor cells was assessed through the MIB-1 antibody that recognizes the Ki-67, being possible to count the cells that are in mitosis.

##### 2.3.2.5 Statistical Analysis

Statistical analysis of “in vivo” studies in nude mice was performed using GraphPad Prism software version 9.0.0. Verification of normality was performed using the non-parametric Shapiro-Wilk test. When normality was verified, an unpaired parametric t-test was subsequently performed, and when this was not verified, a non-parametric test was performed (Mann-Whitney test). All statistical tests were two-sided and *p*-values less than 0.05 were considered statistically significant.

## 3. Results and Discussion

### 3.1 PMC79 targets specifically KRAS hotspot mutations and downstream signaling molecules in CRC

Previous published results from our group revealed that PMC79 possesses a strong anticancer activity in SW480 cell line with KRAS^G12V^ mutation, significantly decreasing proliferation, motility and inducing apoptosis^46^. KRAS and KRAS downstream signaling pathways, MAPK-ERK and PI3K-AKT are involved in the regulation of survival, proliferation and motility, phenotypes of utmost importance in CRC carcinogenesis. We decided to further explore the possible mechanism of action of PMC79 starting by studying its effect on the expression of KRAS and KRAS downstream signaling molecules ERK and AKT. The results demonstrated that PMC79 was able to decrease the expression of KRAS, p-ERK, ERK, p-AKT and AKT in SW480 cells harboring KRAS^G12V^ mutation (Figure 1b). In comparison, cisplatin and other three ruthenium compounds structurally similar to PMC79 (PMC78, LCR134, LCR220) were used and no alterations were observed on KRAS and KRAS downstream signaling molecules expression (Figure 1b and Figure S2).

Besides KRAS^G12V^, present in 9% of CRC, there are other hotspot mutations with relevant prevalence, such as KRAS^G12D^ (13%) and KRAS^G13D^ (7%). Therefore, the effect of PMC79 on the expression of KRAS, AKT and ERK proteins was assessed in CRC cell lines with KRAS^G12D^ and KRAS^G13D^ mutations, respectively LS174T and HCT116 cells. Furthermore, to validate the results obtained for KRAS^G12V^ hotspot mutation in SW480 cells, an additional CRC cell line (SW620) bearing the same mutation was used. The results showed that PMC79 also decreases the expression of KRAS, p-ERK, ERK, p-AKT and AKT in SW620 (KRAS^G12V^) and LS174T (KRAS^G12D^) cell lines (Figure 1c and 1d and Figure S3). In contrast, no alterations were observed in HCT116 cells with KRAS^G13D^ mutation (Figure 1e and Figure S3).

In addition, to disclose the selectivity of PMC79 for mutated KRAS, we used RKO a CRC cell line with KRAS^wt^ protein, and no effect on the expression of KRAS and KRAS downstream proteins was observed (Figure 1f).

Our results suggest that PMC79 might be specific and selectively target KRAS^G12V^ and KRAS^G12D^ mutated proteins and downstream signaling pathways not affecting KRAS wild-type protein and signaling. Moreover, we could observe that PMC79 not only decreases the expression levels of KRAS and activated (phosphorylated) p-AKT and p-ERK but also of the total AKT and total ERK. These results are in accordance with the studies conducted with the first direct KRAS^G12C^ inhibitor recognized as a potential drug candidate, ARS-853, that also showed to reduce KRAS and both phosphorylated and total levels of AKT and ERK proteins in H358 NSCLC-derived cell line with KRAS^G12C^ mutation, after long periods of exposure to the compound^27^.

In the case of specific ERK inhibitors, Balmanno *et al.* also showed that BVD-523, an ERK1/2 inhibitor, decreased p-ERK and total ERK2 expression levels^82^. In a different study, Choi *et al.* demonstrated that the cotreatment of cancer cells with MK-2206 (an inhibitor of AKT activation) and salinomycin, reduced both p-AKT and total AKT levels and sensitizes cancer cells for therapies targeting the PI3K/AKT/mTOR pathway^83^. Therefore, this is not the first time that a compound affects both phosphorylated and total levels of ERK and AKT. Concerning sotorasib and adagrasib it is known that they covalently bind to KRAS^G12C^ resulting in an upward electrophoretic mobility shift of the protein band migration by immunoblot. As consequence of KRAS^G12C^ inhibition, these compounds impair the viability of KRAS^G12C^ mutant cell lines of NSCLC and PDAC, decrease proliferation, induce apoptosis and decrease p-ERK protein levels. In addition, they also inhibit the levels of KRAS-GTP^16,18^. However, neither sotorasib nor adagrasib affect total KRAS expression levels and PI3K signaling. The absence of inhibition of the PI3K signaling pathway might provide an explanation for the acquisition of resistance to these KRAS^G12C^ inhibitors^18^.

It is known that KRAS is located and binds to the inner leaflet of the plasma membrane and its activation occurs at the membrane due to the conversion of GDP to GTP^41^. The presence of a mutation in the kinase domain of KRAS results in KRAS “locking” in the active GTP-bound state resulting in a constitutively activated state at the cell^14^. Previous results from our group showed that PMC79 is preferentially distributed in the membrane fraction^46^ and thus we might hypothesize that this localization might explain in part PMC79 selectivity and inhibitory effect for mutated KRAS.

In order to understand if PMC79 inhibits KRAS by interfering with KRAS activation, we performed an Active Ras Pull-Down assay to determine the RAS-GTP abundance in SW480^KRASG12V^ cells treated with PMC79. We showed that PMC79 reduced the activation of both RAS and KRAS proteins in SW480 cell line with KRAS^G12V^ mutation (Figure 1g).

The major goal of any cancer therapy is to specifically target a protein that is only present in cancer cells and not in normal cells and at the same time interfere with the signaling pathways regulated by them. Inhibition of only MAPK or PI3K pathways is frequently associated with acquisition of resistance to these types of treatment in NSCLC, PDAC and CRC^84,85^. In these cases, it is necessary the combination of different drugs to overthrow this problem^18^. We discovered a novel compound PMC79 that inhibits specifically the expression of KRAS^G12V^ and KRAS^G12D^ mutations not interfering with KRAS^wt^ and also blocks the downstream signaling molecules ERK and AKT both essential in colorectal carcinogenesis. To the best of our knowledge, until now nothing was known about the interaction of ruthenium-derived compounds with KRAS and KRAS signaling pathways^40^. This is the first organometallic ruthenium compound discovered that affects KRAS kinase activation and protein levels as well as both ERK and AKT signaling pathways activation and expression in CRC harboring KRAS mutations. PMC79 has the advantage of specifically inhibiting not only mutated KRAS but also its downstream regulators MAPK and PI3K pathways in CRC cells with KRAS^G12V^ and KRAS^G12D^ mutations. Our new drug PMC79 might constitute a promising novel therapeutic approach to overcome resistance in these types of cancers.

### 3.2 PMC79 is more potent in inhibiting KRAS mutation compared with other KRAS inhibitors

To understand how potent PMC79 was in inhibiting KRAS mutations and signaling pathways it was important to compare the PMC79 effect with other KRAS inhibitors commercially and clinically available. As mentioned above, a series of strategies were developed to indirectly or directly target KRAS in cancer cells^14,86^. The inhibition of KRAS using specific siRNA has been studied as an alternative approach to target KRAS mutations in CRC. Recently, two direct KRAS^G12C^ inhibitors, namely sotorasib and adagrasib were developed and are already in use in the clinic for NSCLC therapy^16,17^.

In this study, we use a siRNA for KRAS and the two KRAS^G12C^ inhibitors adagrasib and sotorasib. We showed that PMC79 inhibits the expression levels of KRAS to a similar extent when compared with the specific siRNA for KRAS (Figure 2a). In addition, no alterations in the expression levels of KRAS were observed in the conditions incubated with adagrasib and sotorasib (Figure 2b).

**Figure 2.**
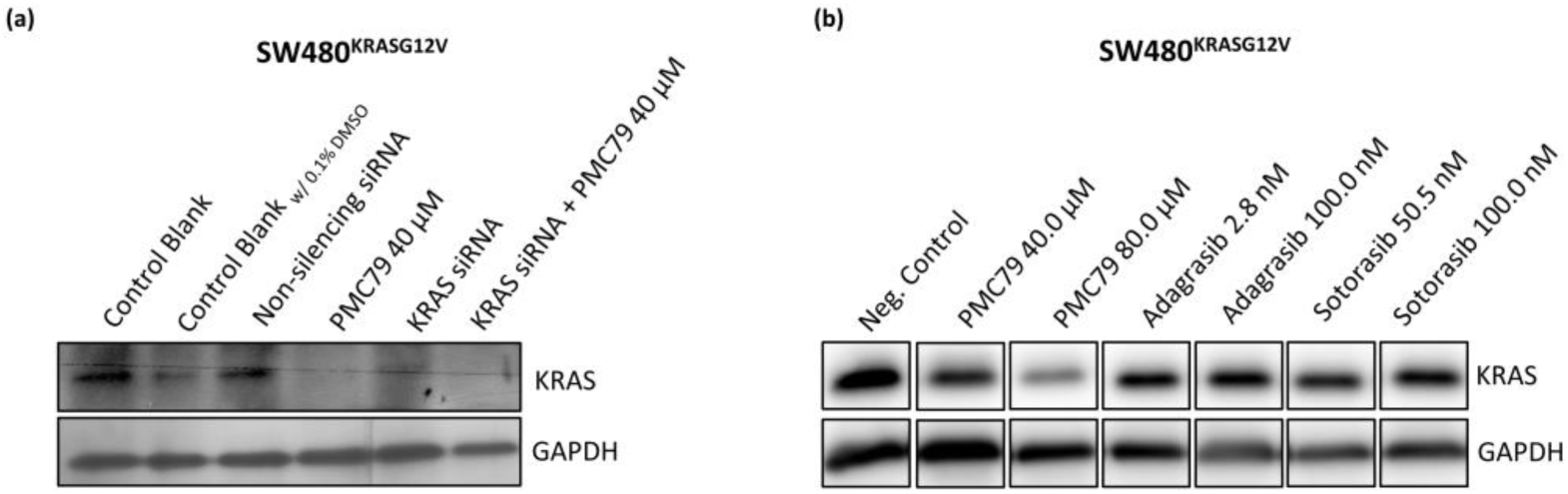
PMC79 KRAS inhibitory effect compared with siRNA of KRAS and adagrasib and sotorasib. **(a)** Immunoblot analysis of KRAS in SW480 cells. Cells were left non transfected (controls), transfected with siRNA control and siRNA targeted against KRAS. After 6 h of transfection, cells were incubated with PMC79 or maintained in complete medium or complete medium with 0.1% DMSO, for 48 h. The results were obtained from two independent experiments. **(b)** Immunoblot analysis of KRAS in SW480 cells after 48 h treatment with PMC79, adagrasib and sotorasib. The results were obtained from at least three independent experiments.

Although specific siRNA for mutated KRAS seem to be very efficient there is a difficulty in finding efficient delivery systems that increase cellular uptake and reduce off-target effects of RNAi^87^. Preclinical efforts took advantage of nanoparticle-based platforms to deliver siRNA. One example is exosomes loaded with specific siRNAs to downregulate the production of mutant KRAS and this approach is under clinical investigation for the treatment of PDAC with KRAS^G12D^ mutation^88^.

The results with adagrasib and sotorasib are in accordance with other published studies performed “in vitro” and “in vivo” that tested these compounds in several cancer cell lines with different KRAS hotspot mutations having only inhibitory effect in KRAS^G12C^ mutant cell lines^15–20^. As mentioned above, preclinical studies have shown that sotorasib and adagrasib selectively inhibit KRAS^G12C^ and decrease the levels of p-ERK protein but not PI3K pathway. These effects were only observed in PDAC and NSCLC cell lines harboring KRAS^G12C^ mutation and not in other cancer cell lines like CRC harboring other KRAS mutations like KRAS^G12V^ and KRAS^G12D 15–20^.

Very recently, other molecules that inhibit hotspot mutations prevalent in CRC are currently in preclinical trials, such as MRTX1133, JAB-22000, RMC-9805 which are direct KRAS^G12D^ inhibitors, JAB-23000 is being studied for KRAS^G12V^ mutations and the JAB-23400 is a multi-KRAS inhibitor. The reason we did not test some of these compounds is because they are not commercially available with the exception of MRTX1133. When we performed this experiment only sotorasib and adagrasib were commercially available. Despite these inhibitors are being tested for possible use in CRC, only KRAS^G12C^ inhibitors^89^ sotorasib and adagrasib are being used at the clinics but only in NSCLC and not in CRC.

### 3.3 PMC79 inhibits KRAS expression in a KRAS-humanized yeast model

To confirm the specific effect of PMC79 in different KRAS hotspot mutations we used a KRAS-humanized yeast model a “clean” cell system using a simple eukaryotic organism^59^. “Humanized-yeast models” emerged as valuable tools for studying human biology. Given the huge number of conserved genes and expression mechanisms similar to humans, simple genome structure, and ease of genetic manipulation these models are valuable for the study of human proteins and drug screening^90^.

KRAS-humanized yeast model using *Saccharomyces cerevisiae* is a well-established model studied in our group for several years^59^. The model’s “cleanness” allows the study of different KRAS hotspot mutations (KRAS^G12V^, KRAS^G12D^ and KRAS^G13D^) in the same genetic background.

We assessed the effects of PMC79 on yeast viability and KRAS expression levels in yeasts with main CRC hotspot mutations and wild-type protein. PMC78, a ruthenium compound derived from PMC79, and adagrasib, a well-known KRAS^G12C^ inhibitor, were also included in these experiments as comparison.

The results show a drastic decrease in viability at the highest dose of PMC79 and PMC78 compounds as early as the first hour of incubation in yeast with mutated KRAS (Figure 3a and Table S2-S6). However, while PMC79 maintains the decrease in viability until 24 h, the effects of PMC78 are recovered over time. At low doses, PMC79 shows little effect on yeast viability, while the highest concentrations of PMC78 show similar results regardless of the dose (Figure 3a and Table S2-S6). On the other hand, adagrasib causes small changes in cell viability in all yeast strains.

**Figure 3.**
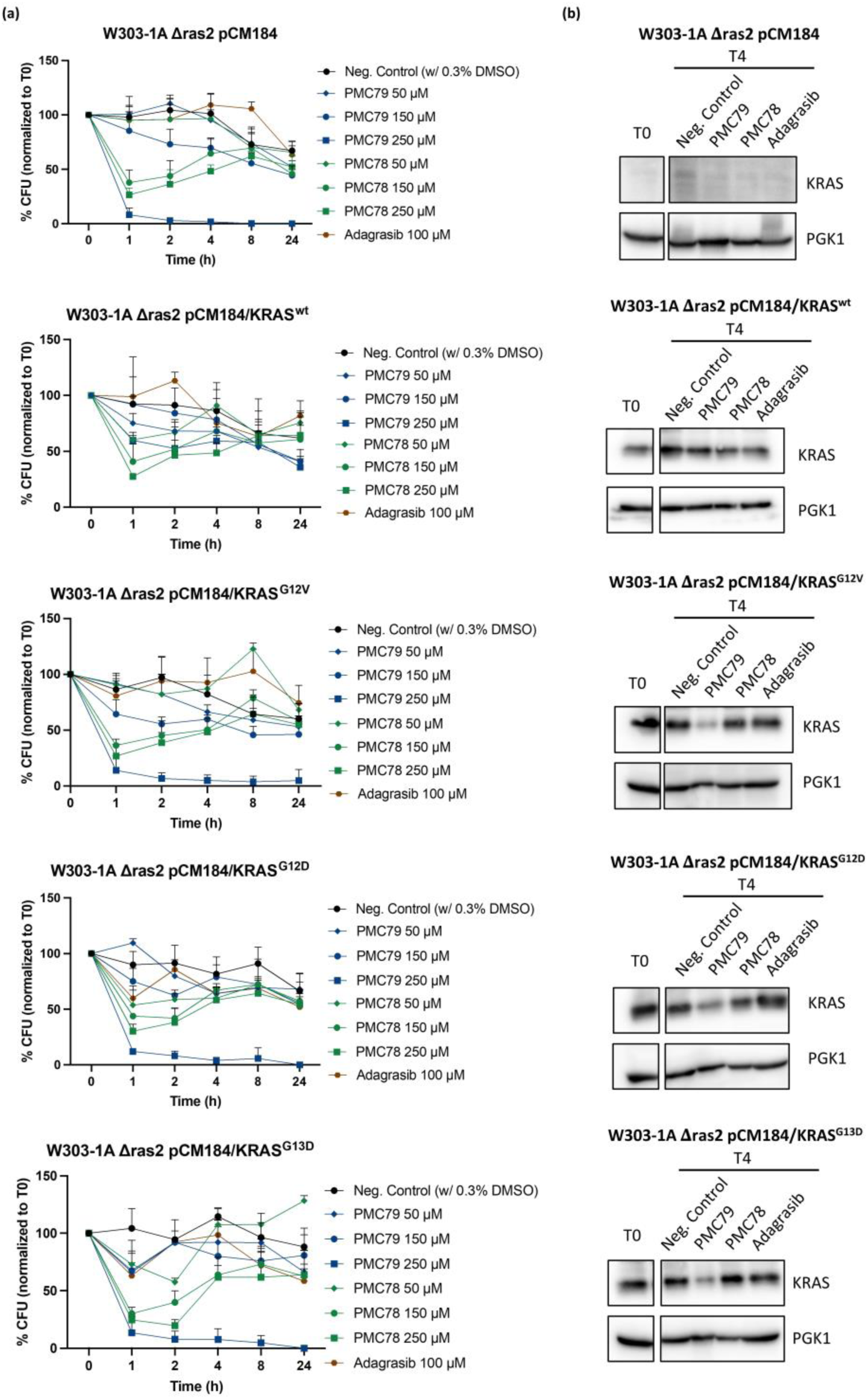
PMC79 also decreases KRAS expression in the KRAS-humanized yeast model. **(a)** Cellular viability analysis of KRAS-humanized yeasts with different CRC hotspot mutations and KRAS^wt^, exposed to several concentrations of PMC79, PMC78 and adagrasib. Data are presented as mean ± SD from at least three independent experiments. **(b)** Immunoblot analysis of KRAS expression in the KRAS-humanized yeast model with different CRC hotspot mutations and KRAS^wt^ after 4 h treatment with PMC79 at 250 µM, PMC78 at 250 µM and adagrasib at 100 µM. The results were obtained from two independent experiments.

Importantly, the results of PMC79 in yeast with mutated KRAS are similar between the different hotspot mutations. Furthermore, the drastic decrease in viability is comparable to that observed in yeast with an empty plasmid (without KRAS) indicating that this compound may target mutated KRAS leading to a phenotype similar to the absence of this protein. In yeast with KRAS^wt^, PMC79 decrease in viability is not pronounced, indicating that the presence of the normal protein confers resistance to the compound. In contrast, PMC78 and adagrasib are not able to differentiate between mutated and wild-type KRAS proteins (Figure 3a and Table S2-S6).

The next step was to study the KRAS expression levels at 4 h of incubation with the highest doses of the three compounds. Similar to what was observed previously in CRC-derived cell lines, PMC79 also decreased KRAS expression in yeast with mutated KRAS but did not alter the expression levels of KRAS^wt^ (Figure 3b). In contrast, PMC78 and adagrasib do not affect the expression of KRAS^G12V^, KRAS^G12D^, KRAS^G13D^ and KRAS^wt^. On the other hand, in comparison with adagrasib, PMC79 seems to be more specific to target the main hotspot mutations found in CRC.

Contrary to what was observed in the HCT116^KRASG13D^ CRC cell line, in yeast, PMC79 also showed to affect the yeast strain containing KRAS^G13D^ mutation and decrease the expression levels of this hotspot mutation. This result may be due to the fact that HCT116 cell line has a complex genetic background with other mutations besides KRAS, namely PI3K mutation. Indeed, both PI3K and MAPK signaling pathways are known to co-regulate each other and upstream KRAS and this might result in resistance to PMC79^91^.

KRAS-humanized yeast model is a simple and “clean” model and the results obtained showed that PMC79 decreases the viability of yeasts expressing hotspot KRAS mutations and not in yeast expressing KRAS^wt^ but only inhibiting KRAS expression levels in cells harboring those mutations. This result also proves that the KRAS-humanized yeast model is an excellent tool for studying the effect of new drugs on KRAS mutations.

Overall, PMC79 results in the CRC-derived cell lines and in the KRAS-humanized yeast model suggest that PMC79 is selective for mutated KRAS. In the case of yeast strains expressing KRAS^wt^ protein, the cells seem to be more resistant to PMC79 and do not affect KRAS^wt^ protein expression. These results suggest that the mechanism of action of PMC79 might be related to the structural conformation of the mutated protein that might be more available for the binding of the compound and thus allow the inhibition of the mutated protein.

Taking into consideration the effect of PMC79 on the specific inhibition of all KRAS hotspot mutations, we decided to gather evidence on the mechanism by which PMC79 might inhibit KRAS mutation activation and expression levels.

### 3.4 Molecular docking results

In an attempt to gather insights into the binding of PMC79 to wild-type KRAS (WT) as well as G12X mutations (G12D, G12V), we used a combination of MD simulations and molecular docking to hopefully build a tentative structural model for the interaction. We explored possible binding near the P-loop region, responsible for binding phosphate and regions near Switch-I and Switch-II regions that are not only responsible for KRas/SOS1 and K-Ras/NF1 interactions^92^ but also provide a shallow pocket for small-molecule binding^93^. These switch regions are typically disordered in crystallographic structures and hence, the usage of X-ray structures as receptors for docking studies would be pernicious.

To minimize the bias of the crystallographic structure and provide sampling of these regions, we docked compound PMC79 into 180 conformations of each receptor (WT, G12D, and G12V) and selected the lowest-energy binding poses according to Vina and Vinardo scoring functions for further analysis. These are represented in Figure 4. Apart from the WT, it appears to exist a relatively good consensus in the binding mode provided by both scoring functions. Indeed, for G12D, PMC79 is located below Switch-II, with the cyclopentadienyl ligand pointing to the protein exterior while the phenyl groups of the triphenylphosphine establish hydrophobic interaction with Switch-II. The polar hydroxyl groups are hydrogen bonded to the side-chain of ASP69 (Vina) and eventually to the main-chain oxygen and nitrogen atoms of GLN99 (Vinardo). No interaction with ASP12 was observed, despite the presence of the polar hydroxyl groups on the bipyridine ligand. In the lowest-energy binding poses of G12V, PMC79 binds differently as the compound is now under Switch-II but with the one hydroxyl group establishing hydrogen bonds with the main-chain oxygen of VAL8 (Vina and Vinardo) near the P-loop, in a much close proximity to the mutated residue of interest (VAL12), though no specific interaction is observed with the latter. This close interaction under Switch-II along with a close proximity with the P-loop might explain the larger activity of PMC79 against this mutant.

**Figure 4.**
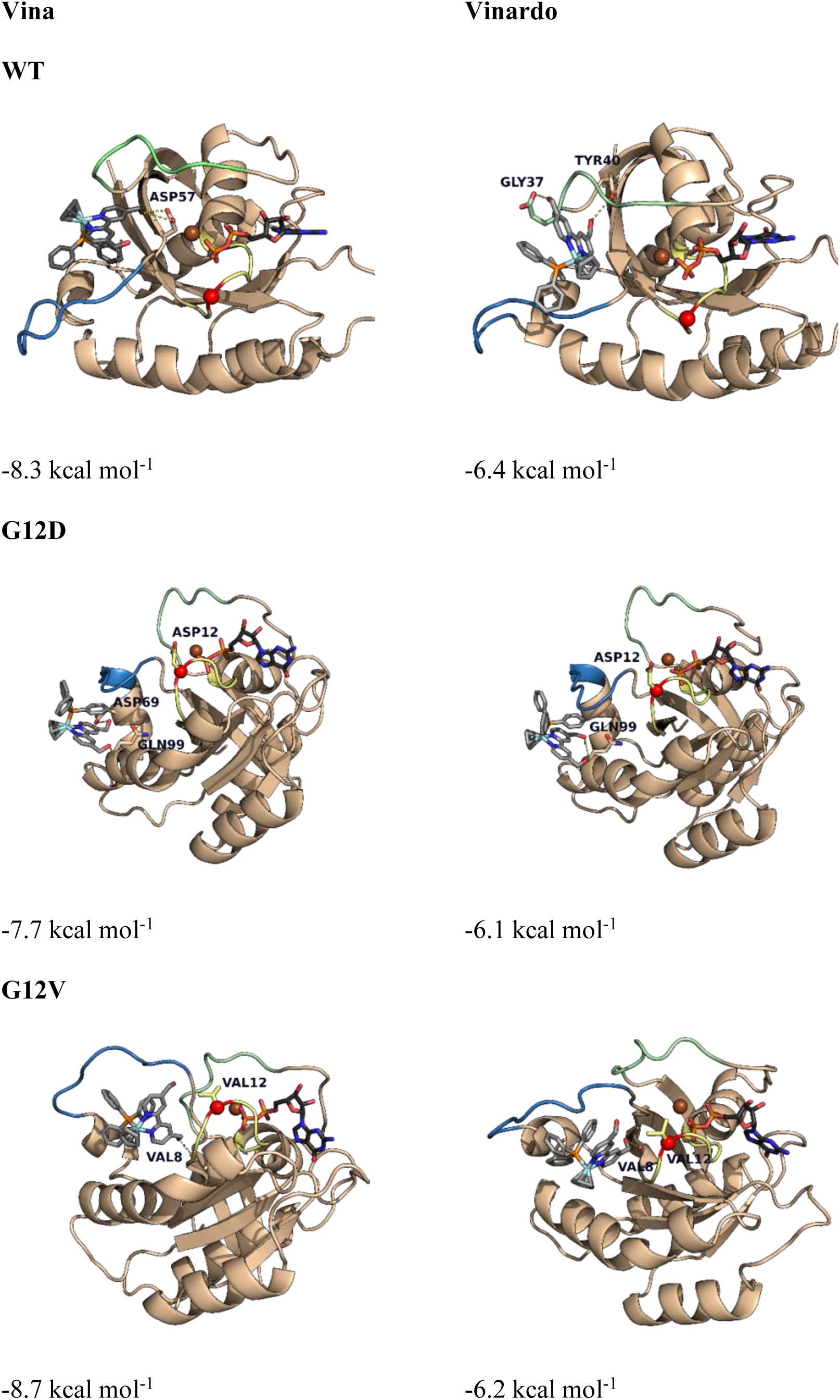
Lowest-energy binding poses obtained with Vina and Vinardo scoring functions for PMC79 docked into KRAS: WT (top), G12D (middle), G12V (bottom). Hydrogen bonds between PMC79 the receptor are represented in yellow dashes. The remainder details as in Figure S1.

For the interaction with the wild-type, consensus was not observed in the binding mode. Vina scoring function indicates that PMC79 sits between both switch regions, with cyclopentadienyl and triphenylphosphine directed towards the protein exterior while hydrogen bonds are established with the side-chain of ASP57. On the other hand, using Vinardo scoring, the compound is also located sits between both switch regions but rotated, with hydrogen bonds with TYR40 and GLY37 (Switch-I).

If one takes into consideration the actual scores of the binding poses, Vina predicts G12V > WT > G12D while for Vinardo the order is WT > G12V > G12D. Both docking scoring functions failed to predict the observed activity though Vina was able to order G12V as the best mutant for PMC79 binding as per experimental results. This difficulty in predicting the correct order is, besides the inherent problems of docking scoring functions^94^, also largely due to the fact that KRAS dynamics is particularly challenging to tackle, and that phenomenon is paramount for ligand binding. In our poses with G12V, the PMC79 ligand is much closer to the GDP site and that is only possible on a sampled protein conformation where the P-loop approaches the Switch-II region in some extent. Indeed, long MD simulations indicated that G12D mutants possess a closer dynamics to the wild-type than G12V^95^. Therefore, our results suggest that the specific activity of PMC79 towards G12V, which is reduced in G12D and absent in the wild-type, is not due to a specific interaction of the compound with the G12X mutation, as this kind of interactions was not found, but rather due to changes in the protein dynamics which are mutation specific and were previously shown to affect allosteric communication networks^95^, eventually enabling the binding in G12V (and to a lesser extent G12D) while hindering the interaction with WT. Such change of the protein dynamics upon mutation was also observed for G13^96^ and Q61 mutants^97^. The complexity of this mechanism deserves further studies, beyond the limitations of molecular docking, but are however out of the scope of this manuscript.

### 3.5 KRAS inhibition by PMC79 is not dependent on actin polymerization or on proteasome

Actin cytoskeleton is known to be responsible for KRAS clustering and localization at the cell membrane through the formation of KRAS nanoclusters with the inner leaflet of the plasma membrane which is important for KRAS signal transduction^98^. Our previous results showed that PMC79 interferes with actin cytoskeleton in CRC cell lines leading to alterations in cell-cell adhesion and intercellular contact establishment, inducing changes in cell phenotype and roundness and reducing the expression levels of ß-actin^46,47^. Therefore, to prove if the inhibition of KRAS expression by PMC79 might be dependent on actin filament’s function, we studied the influence of latrunculin A, an actin polymerization inhibitor on the effect of PMC79. We performed a phalloidin staining and as already described before, PMC79 affects cell-cell adhesion and intercellular contact establishment and induces changes in cell phenotype and roundness^46,47^ (Figure 5a). Concerning latrunculin A it also leads to actin structure alterations consistent with depolymerization of actin and when combined with PMC79 similar actin phenotype was observed, with evident changes in cell phenotype similar to PMC79 alone (Figure 5a).

**Figure 5.**
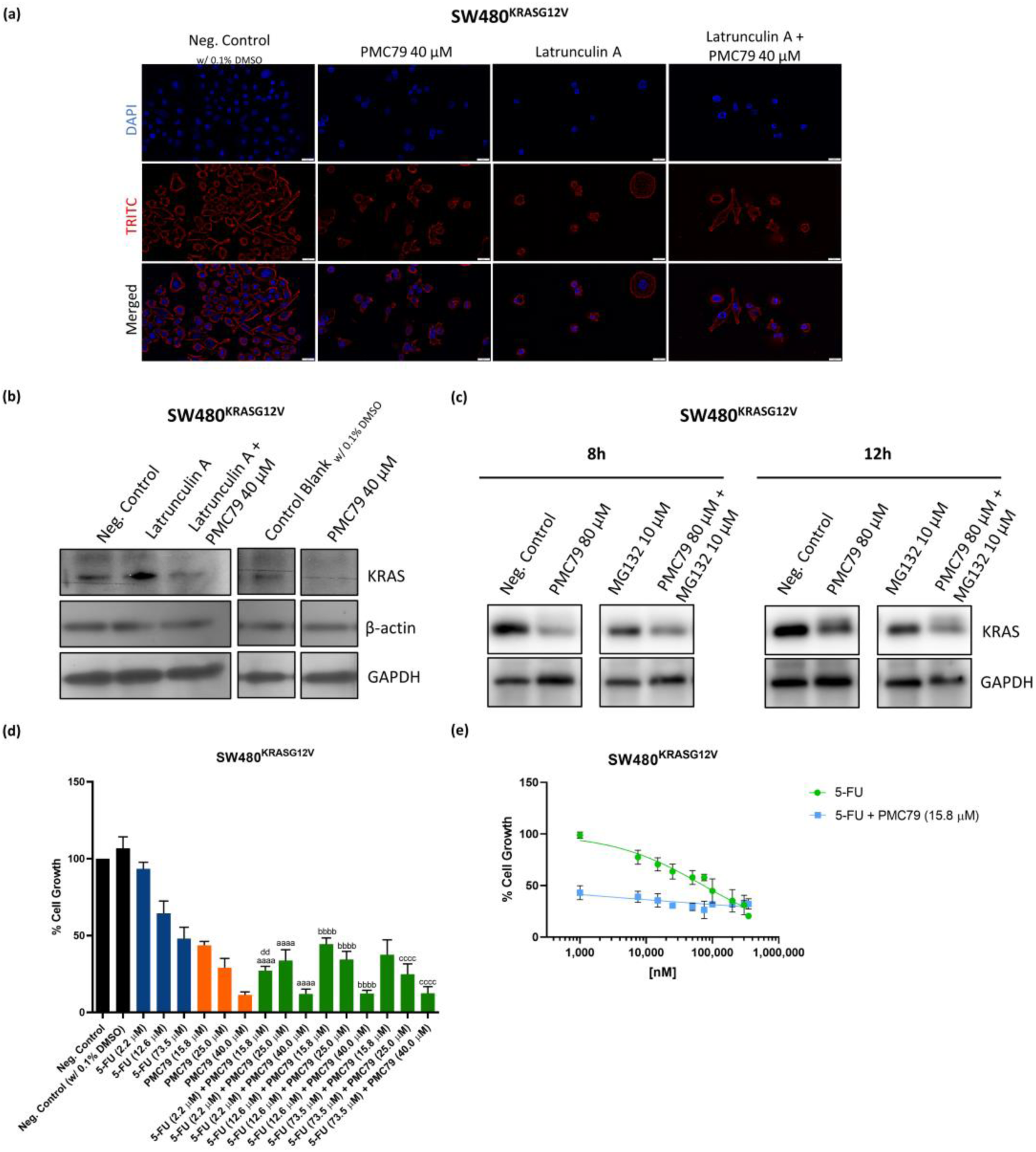
PMC79 mechanism of KRAS inhibition. **(a)** Representative images (×600) of DAPI (4′,6diamidino-2-phenylindole), Phalloidin-AlexaFluor®568, and their merge were obtained with fluorescence microscopy. The results were obtained from at least three independent experiments. Scale bar is 20 μm. **(b)** Immunoblot analysis of KRAS and β-actin in SW480 cells after 48 h treatment with PMC79 and latrunculin A. The results were obtained from two independent experiments. KRAS expression levels of PMC79 and respective control are the same of the figure 2a because experiments were conducted at the same time. **(c)** Immunoblot analysis of KRAS in SW480 cells. Cells were pretreated with MG132 for 1 h and then co-treated with PMC79 for 8 h and 12 h. The results were obtained from at least three independent experiments. **(d)** “In vitro” effects of different doses of PMC79 and 5-FU alone and in combination on cell growth of SW480 cells, determined by SRB assay, after 48 h of treatment. **(e)** Dose-response curves of 5-FU alone and in combination with PMC79 at 15.8 μM. Values represent mean ± SD of at least three independent experiments. ^aaaa^ p ≤ 0.0001 compared with 5-FU (2.2 μM); ^bbbb^ p ≤ 0.0001 compared with 5-FU (12.6 μM); ^cccc^ p ≤ 0.0001 compared with 5-FU (73.5 μM); ^dd^ p ≤ 0.01 compared with PMC79 (15.8 μM).

Our results on the analysis of KRAS expression levels showed that PMC79 inhibits KRAS expression even in the presence of latrunculin A, indicating that KRAS expression inhibition by PMC79 is not an indirect effect that this compound has on the actin cytoskeleton and thus is not dependent on the actin structure at the cell (Figure 5b).

In order to understand the reason why PMC79 leads to a decrease in the expression levels of mutated KRAS, we hypothesized that PMC79 could target mutated KRAS to proteasomal degradation. The proteasome is a multiprotein cellular complex that regulates most proteins and in consequence several cellular processes with high importance for carcinogenesis such as proliferation, apoptosis, angiogenesis and motility/metastasis^99^. CRC is not an exception with several pathways, such as Wnt and KRAS pathways, that play important roles in carcinogenesis being regulated by the proteasome complex^100^. Some studies already reported that KRAS and KRAS downstream signaling molecules are regulated and consequently degraded by proteasome ^101–103^. This strategy is one of the approaches used that can directly target KRAS and recently, the natural product, Kurarinone, showed to reduce KRAS protein levels in CRC cells, through proteasomal degradation dependent on an E3 ubiquitin ligase WDR76^103^.

Thus, to assess whether the inhibition of the ubiquitin-proteasome pathway restores KRAS expression after PMC79 treatment, we used a well-known proteasome inhibitor, MG132. PMC79 effect on the KRAS half-life showed this compound begins to decrease KRAS expression after 8 h of treatment (Figure S4). Therefore, MG132 was pre-incubated with SW480^KRASG12V^ cells for 1 h and then co-incubated with PMC79 for additional 8 h and 12 h. The results showed that even in the presence of MG132, PMC79 keeps decreasing the expression of KRAS, suggesting that KRAS inhibition by PMC79 is independent of proteasomal degradation (Figure 5c).

Summing up, in the present study, we showed that PMC79 inhibits specifically KRAS hotspot G12V and G12D mutations expression and interferes with ERK, AKT phosphorylation and expression in CRC cells. We demonstrated that at least in CRC cells harboring KRAS^G12V^ mutation PMC79 impairs KRAS-GTP binding and consequent activation.

Using the KRAS-humanized yeast model we could also observe that PMC79 might also interfere with KRAS^G13D^ mutation expression. PMC79 does not interfere with KRAS wild-type both in CRC cells or KRAS-humanized yeast model.

In order to try to understand the possible mechanism of action of PMC79 that could explain the specific inhibitory effect of PMC79 on KRAS hotspot mutations and downstream signaling pathways, here we explored some processes implicated in the regulation of protein expression and activation. Molecular docking study did not find any interaction of the compound with mutated KRAS but did not exclude the possibility of some interaction due to changes in the protein dynamics which are mutation specific. The results gathered here suggest that it might be the protein dynamics associated with the mutations that can be responsible for the specific effect of PMC79 in KRAS hotspot mutations. Moreover, our results also suggest that PMC79 seems to inhibit KRAS protein expression levels independently of actin cytoskeleton and proteasome.

### 3.6 PMC79 potentiates 5-FU anticancer effect on SW480^KRASG12V^ cell line

Regardless of the mechanism by which PMC79 inhibits specifically KRAS hotspot mutations in CRC, it is of utmost relevance to study the anticancer effect of PMC79 in CRC harboring KRAS mutations. For that matter, we tested the combination of PMC79 with 5-FU a classical chemotherapeutic drug to compare their anticancer effect and explore the combination of the two drugs as a new therapeutic approach.

5-FU is the conventional chemotherapeutic drug used in the treatment of CRC blocking DNA replication however, it is associated with low success rates, many side effects and resistance problems^104^. Previous results from our group already showed that PMC79 induces several phenotypic alterations in hallmarks of CRC with KRAS mutation, namely decreases proliferation and induces cell cycle arrest, decreases colony formation, induces apoptosis and necrosis and reduces motility of SW480^KRASG12V^ cells^46^.

We tested the effect of different concentrations of PMC79 and 5-FU alone and in combination to study their effect on SW480^KRASG12V^ cells viability. Almost all combinations decrease cell viability compared with the same dose of 5-FU alone (Figure 5d). However, only the combination of the lower dose of the two agents (5-FU 2.2 μM + PMC79 15.8 μM) led to a significant decrease in cell viability regarding the same concentrations of PMC79 and 5-FU alone.

To confirm that the lowest concentration of PMC79 (15.8 μM) potentiates 5-FU anticancer effects, the dose-response curves of 5-FU alone and in combination with PMC79 were determined. The results show that the combination of 5-FU with PMC79 strongly inhibits cell growth at the lowest doses of 5-FU, reducing cell growth by half (Figure 5e). In addition, 5-FU half-maximal inhibitory concentration (IC_50_) reduced drastically in combination with PMC79, from 73.5 μM to 0.02 μM (Table S7).

These promising results indicate that low doses of PMC79 potentiate 5-FU anticancer effects in a CRC cell line with KRAS mutation.

Several studies have shown that combining 5-FU with other therapeutic agents increases treatment success rates in addition to decreasing side effects since lower doses of each agent are required^104^. Our data open the possibility of using PMC79 in combination with 5-FU as a new therapeutic approach in CRC with KRAS mutations targeting specifically mutated KRAS, interfering with ERK, and AKT pathways while blocking DNA replication of these cells.

### 3.7 PMC79 reduces tumor growth in the “in vivo” CRC chick embryo chorioallantoic membrane-xenograft

In order to do an “in vivo” “proof of concept” we performed pre-clinical trials using two different complementary models, the chick embryo CAM model and the CRC-mice xenografted model.

Previous “in vitro” results showed that PMC79 presents good anticancer activity against the CRC cell line SW480 with KRAS^G12V^ mutation^46^ besides inhibiting KRAS expression. Therefore, we evaluated the “in vivo” anticancer effects of PMC79 using the chick embryo CAM-SW480 xenograft assay.

SW480 cells grew in CAM from day 9 to day 13 when tumor xenografts were exposed to different treatments for 4 days. In the end, tumor growth was determined through the area difference between day 13 and day 17. The results showed the highest dose of PMC79 (40.0 μM) showed a decrease in the tumor area and the dose of 73.5 μM of 5-FU demonstrated a tendency towards a decrease in the tumor area. The lower doses of PMC79 (25.0 μM) and of 5-FU (12.6 μM) have no effect on the tumor area (Figures 6a and 6b).

**Figure 6.**
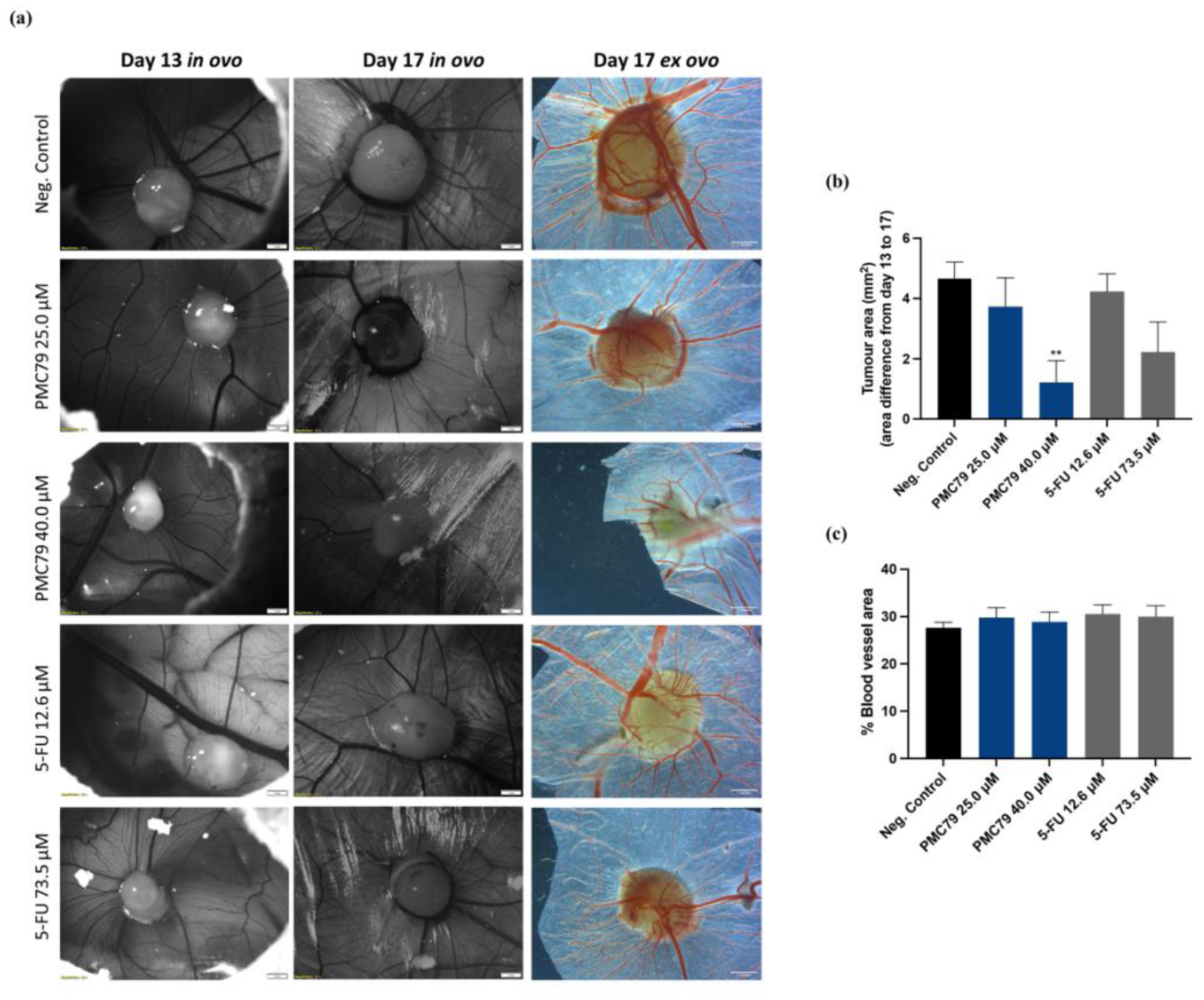
PMC79 reduces tumor growth in the chick embryo CAM assay. **(a)** Representative images of “in vivo” CAM assay “in ovo” and “ex ovo” before (day 13) and after treatment (day 17). **(b)** Tumor area analysis (area difference between day 13 and day 17) of “in vivo” CAM assay. **(c)** Graphical representation of the percentage of blood vessel formation of “in vivo” CAM assay. Results are expressed as mean ± SEM of at least 10 eggs per experimental condition. ** p ≤ 0.01 compared with negative control. Scale bar is 1 mm to “in ovo” images, 1 inches to “ex ovo” images.

Concerning the percentage of blood vessel area around the tumor no alterations were observed for any of the drugs (Figure 6c). These results showed that PMC79 reduces significantly the tumor area in the chick embryo CAM assay not affecting angiogenesis.

The chick embryo CAM-xenograft is a cost-effective and valuable alternative method to study the anticancer effect of drugs with several advantages over the xenograft mice model such as easy access for manipulation, shorter experimental times, physiological environment and reproducibility, and natural immunodeficient model which accept cancer cells regardless of their origin without immune response^105,106^. This data on PMC79 is a preliminary “proof of concept” that the compound might have a valuable anticancer effect by reducing the xenograft of CRC in chick embryo CAM “in vivo”.

### 3.8 PMC79 anticancer effect in the “in vivo” colorectal cancer xenograft mice model

This study aimed to use an “in vivo” xenograft mice model to corroborate the “in vivo” results on chick embryo CAM-CRC xenograft and also to confirm “in vivo” the effect of PMC79 in significantly decreasing KRAS^G12V^ expression in SW480 cells. For that matter, we used N:NIH(S)II-nu/nu mice in which SW480 cells harboring KRAS^G12V^ mutation were injected in the back of the mice in order to produce tumor xenografts.

#### 3.8.1 Toxicity assay

To set the maximum tolerated dose of PMC79 in N:NIH(S)II-nu/nu mice, we first performed a dose escalation experiment. For this, the following doses of PMC79 were tested: 34 mg/Kg, 26 mg/Kg, 17 mg/Kg, 13 mg/Kg and 8.7 mg/Kg (n=4 mice per group, except in the last one where n=2). The 34 mg/Kg dose was found to be lethal. The 26 mg/Kg dose is not advised because it caused 75% of lethality after the first injection. In general, the injection of PMC79 causes abdominal discomfort and agitation followed by apathy, being these effects dose-dependent. Body weight loss was also found to be dose-dependent. Despite these effects, a drug habitation of the animals to PMC79 seems to occur because after several injections the reaction of the animals is not as severe as in the first injection and their weight also increases (Figure 7a). The 17 mg/Kg dose was established as well tolerated by the animals and as such, this dose was selected to be used in the evaluation of the antitumor ability of PMC79.

**Figure 7.**
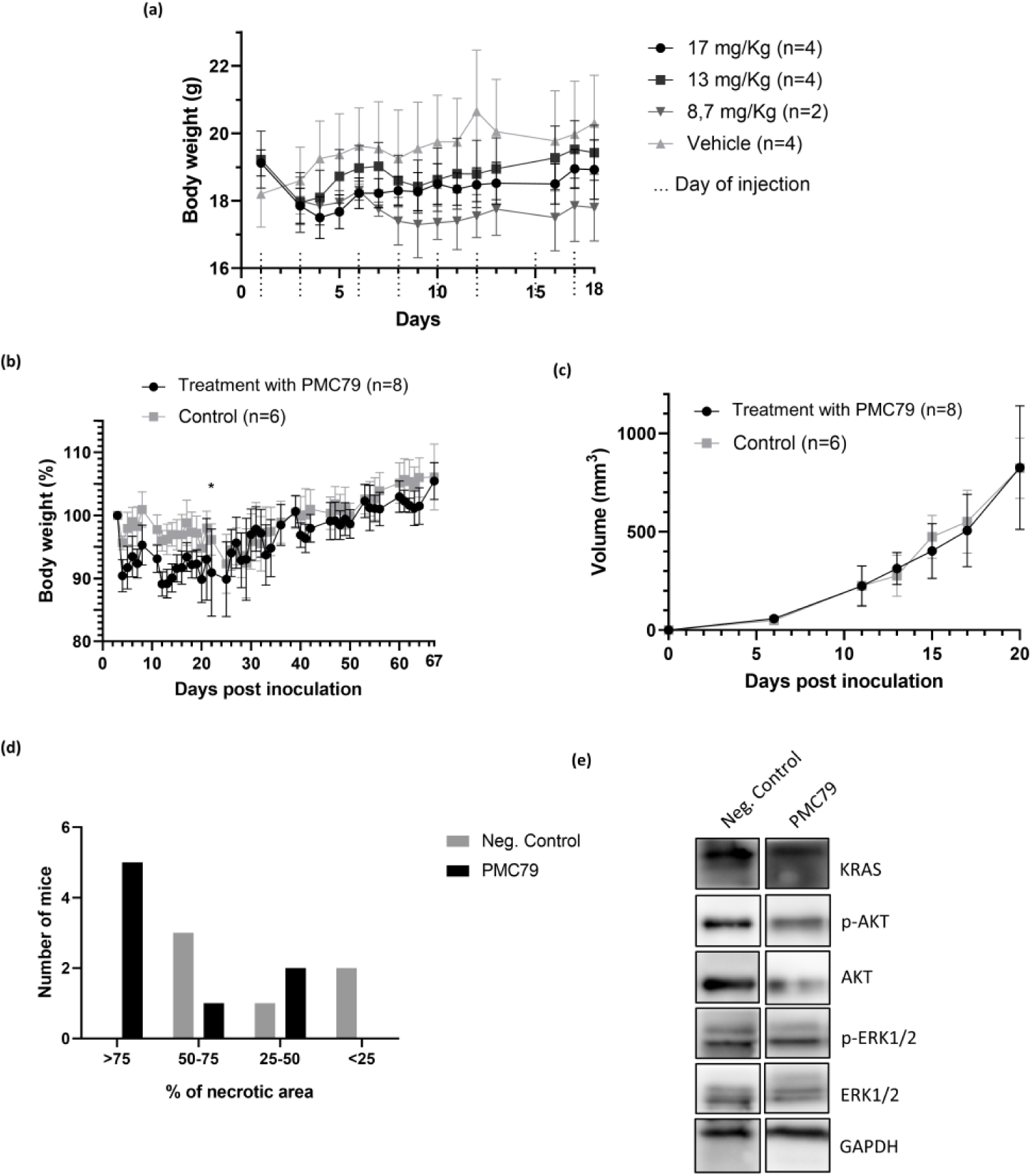
PMC79 antitumor effects in a xenografted CRC mice model. **(a)** Effect of PMC79 in animal wellbeing, measured in terms of body weight. The number of animals used for each group is indicated in the figure. Animals that received the 8.7 mg/Kg dose only began to be injected on day 3. **(b)** Weight evolution of female nude N:NIH(S)II-nu/nu mice treated intraperitoneally with 17 mg/Kg of PMC79 with 40% Captisol® in Milli-Q water (m/v) and vehicle. The number of mice used for each group is indicated in the figure. * From day 22 the PMC79 treatment group consists of only 7 mice due to a sudden weight loss of one mouse due to an infection **(c)** Effect of PMC79 in tumor volume evolution resulting from the subcutaneous inoculation of 1.0×10^6^ SW480 cells inoculated subcutaneously in N:NIH(S)II-nu/nu nude female mice. The number of mice used for each group is indicated in the figure. At day 20 post-inoculation started the excision of tumors. **(d)** Graphic representing the percentage (%) of necrotic area of the entire tumor area (percentage indicated only by observing the tumors under the microscope by the pathologists). **(e)** Results obtained from Western blot analysis to evaluate the expression of KRAS, p-AKT, AKT, p-ERK, and ERK proteins.

From the histological analysis, it was possible to verify that there are two mainly relevant effects that can be associated with the toxicity of the compound (Figure S5). Of the 10 mice administered with PMC79, 5 of them had hemorrhaged lungs, with a correlation between the development of pulmonary hemorrhage and the administered dose of PMC79. In contrast, no bleeding was observed in control mice. These hemorrhages can be derived from a pulmonary alteration that is caused by the administration of PMC79 which can cause the destruction of vessels associated with the spasms/shock situations that the animals present right after the injection. Beyond the hemorrhaged lungs, all animals, including controls showed peripancreatic inflammatory infiltrate, a normal reaction resulting from the IP injections. There was a tendency for more severe peritonitis in mice treated with PMC79 and this toxicity correlates with the administered dose.

To further confirm PMC79 safety, we measured several hematochemical parameters (urea, creatinine, alanine aminotransferase (ALT) and aspartate aminotransferase (AST)) at the end of the study (Table S8). No significant changes were observed between the treated and control Groups. As such we can conclude that PMC79 did not induce biochemical changes in the blood, at least for these biochemical markers.

#### 3.8.2. “In vivo” antitumor activity in a CRC xenograft mice model

N:NIH(S)II-nu/nu mice with SW480 cells tumor xenografts were treated with PMC79 at the established dose of 17 mg/Kg. As shown in Figure 7b, a treatment dose of 17 mg/Kg PMC79 or vehicle leads to weight loss after the first treatment, yet afterward the tendency is to increase weight. Indeed, all animals recovered or increased their initial weight in relation to the first day of treatment.

Concerning the effect of PMC79 in tumor growth, on the conditions of this study no differences were found between tumor volumes of PMC79 treated and control groups (Figure 7c).

After the excision of the tumors, a sample was fixed in 10% buffered formalin and a histopathological and immunohistochemical analysis was carried out. Tumor samples were compared with those of control mice and the histological variables analyzed, namely necrosis, intratumoral hemorrhage, lymphocytic infiltration, peritumoral oedema, as well as the degree of tumor regression were evaluated. The number of mitoses was quantified as well. In general, there are no differences between the group treated with PMC79 and its control, being the cellular proliferative index Ki-67 very high in all cases (data not shown). Interestingly the percentage of necrotic area within the tumor is much higher in tumors treated with PMC79 compared with untreated controls (Figure 7d).

Although the percentage of necrosis for each tumor is merely indicative, the difference between treated and control tumors is remarkable as the animals treated with PMC79 presented tumors with a larger necrotic area compared to the controls. This result is in fact interesting because, although there were no differences in tumor growth between treated and control groups, PMC79 caused the death of tumor cells. This is a fact that corroborates the results obtained *in vitro* in which it was observed cell death of SW480 cells by apoptosis and also necrosis when exposed to PMC79^46^.

In order to infer about possible cancer recurrence and/or the appearance of metastases, we surgically removed the tumors and kept the mice until 67 days after tumors inoculations and 35 days after removal of the last tumors. At this date, all animals were apparently in good health. When carrying out the necropsy of the animals treated with PMC79 and controls, lungs and lymph nodes (4 submandibular, 4 axillary and 2 inguinal) were collected, to search for micrometastases. After histological analysis, it was possible to verify that none of the mice had evident signs of metastatic invasion in the organs analyzed.

#### 3.9.3. Western blot analysis of KRAS signaling pathways

After the excision of the tumors, we performed protein extraction and analyzed the expression of KRAS and downstream signaling pathways AKT and ERK proteins by Western blot. As shown in Figure 7e, PMC79 was able to inhibit the expression of KRAS, ERK, p-ERK, AKT and p-AKT proteins. The results in the “in vivo” mice model-CRC xenograft corroborated the “in vitro” results in SW480^KRASG12V^ cells showing that PMC79 “in vivo” was also able to inhibit the expression levels of mutated KRAS and the downstream proteins ERK and AKT. Overall, these results highlight the therapeutic potential of PMC79 for the treatment of patients with CRC harboring KRAS mutations inhibiting at the same time KRAS, and ERK and AKT signaling molecules.

## 4. Conclusion

KRAS was considered “undruggable” for a long time. After decades of efforts, this paradigm has begun to change with several inhibitors being discovered and two already being used in the clinic for NSCLC treatment. The difficulties in finding new agents that inhibit this molecule are related to its conformational dynamics that are different from mutation to mutation, which limits the application in different KRAS hotspot mutations. KRAS^G12C^ inhibitors have been the most explored so far. However, this mutation is only present in 3% of CRCs with KRAS mutation. KRAS-activating mutations are the most frequent oncogenic alterations in CRC, but the therapeutic options in these types of cancers are scarce^3^. To the best of our knowledge, so far, no KRAS inhibitors for CRC treatment have been accepted in the clinics.

In this work, we discovered for the first time a new ruthenium-derived organometallic agent, PMC79, able to inhibit specifically the expression of various KRAS hotspot mutations in several CRC-derived cells, in KRAS mutated-humanized yeast model and in CRC-xenograft mice “in vivo” model. Moreover, PMC79 also inhibits the expression of KRAS downstream signaling molecules ERK and AKT in CRC cells harboring hotspot mutations. Kinase activity results in CRC cells also demonstrated that PMC79 inhibits KRAS activation. The molecular docking study suggests that the protein dynamics associated with the mutations might be responsible for the specific effect of PMC79 in KRAS hotspot mutations. We also demonstrated that KRAS inhibition by PMC79 is not dependent on actin polymerization or on the proteasome. Importantly, we showed that low doses of PMC79 potentiate 5-FU anticancer effects in a CRC cell line with KRAS mutation, which open the possibility of using PMC79 in combination with 5-FU as a new therapeutic approach in CRC with KRAS mutations.

Here, we performed a “proof of concept” for the anticancer effect of PMC79 “in vivo” showing that in the chick embryo CAM-xenograft model, PMC79 reduced tumor growth and in the xenograft mice model PMC79 induced necrosis of the tumor and decreased the expression levels of KRAS, AKT and ERK proteins.

Taken together, our findings highlight the discovery of the first Ru-derived organometallic compound, PMC79, as a specific KRAS mutated inhibitor in CRC with KRAS mutations. This discovery opens the possibility to study the effects of the compound in other human cancer types also presenting these KRAS mutations.

In this work, we discovered a new compound, PMC79, that specifically inhibits KRAS and both KRAS signaling pathways (MAPK and PI3K) in cells harboring KRAS mutation but not in cells with KRAS wild-type. This simultaneous inhibition of MAPK and PI3K pathways is an advantage over the above inhibitors and may prevent potential resistance problems in the future.

Overall, the results gathered here show a promising therapeutic application of PMC79, supporting the use of this compound as a new specific and potent KRAS signaling inhibitor agent in CRC harboring KRAS hotspot mutations.

## Acknowledgments

We thank Joana Silva for her help on “*in vivo”* protein extraction.

