## Supplementary Figures for "Shifting KRAS hotspot mutations inhibition paradigm in colorectal cancer"

### Slide 1
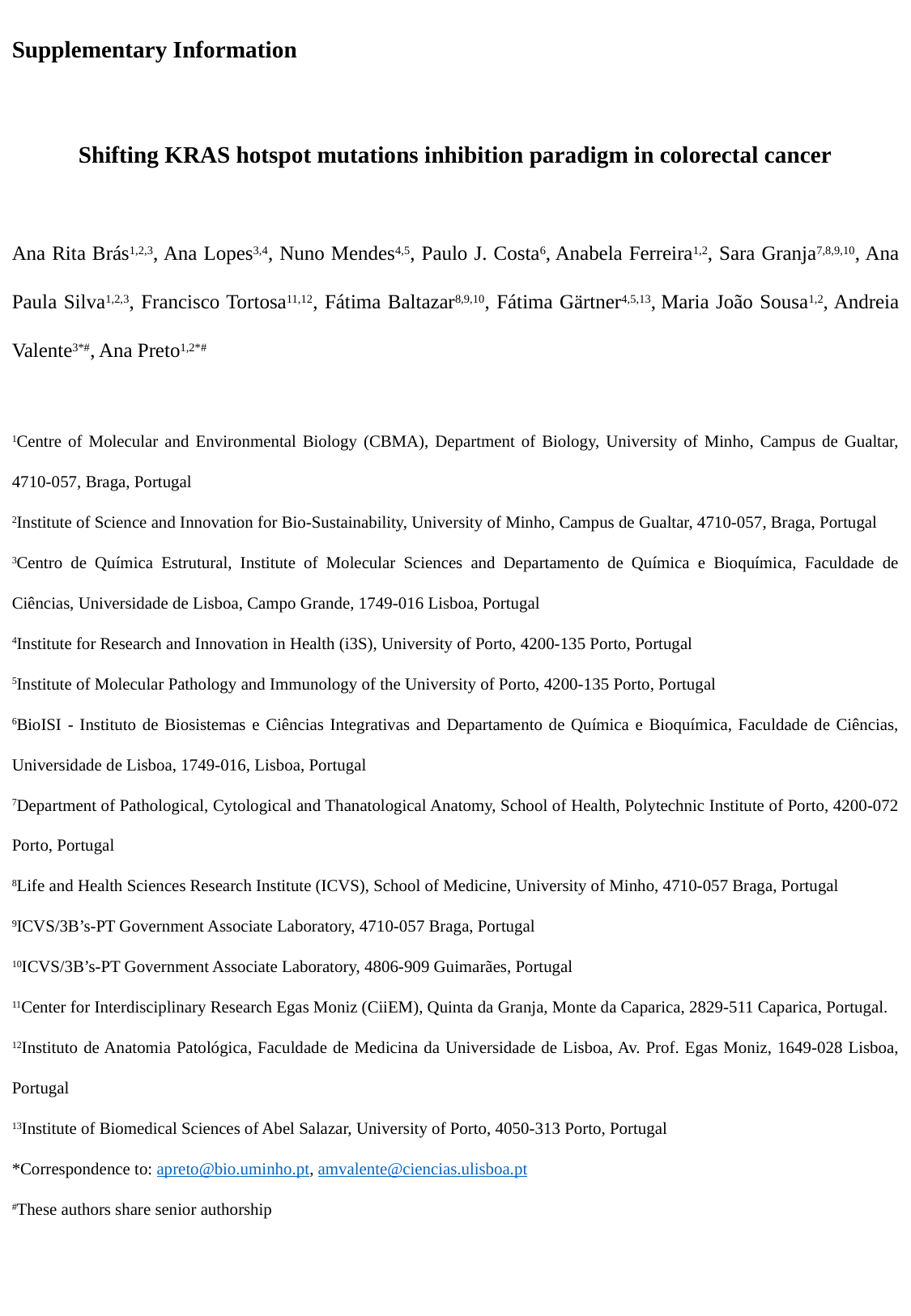

Supplementary Information
Shifting KRAS hotspot mutations inhibition paradigm in colorectal cancer
Ana Rita Brás^1,2,3^, Ana Lopes^3,4^, Nuno Mendes^4,5^, Paulo J. Costa^6^, Anabela Ferreira^1,2^, Sara Granja^7,8,9,10^, Ana Paula Silva^1,2,3^, Francisco Tortosa^11,12^, Fátima Baltazar^8,9,10^, Fátima Gärtner^4,5,13^, Maria João Sousa^1,2^, Andreia Valente^3*#^, Ana Preto^1,2*#^
^1^Centre of Molecular and Environmental Biology (CBMA), Department of Biology, University of Minho, Campus de Gualtar, 4710-057, Braga, Portugal
^2^Institute of Science and Innovation for Bio-Sustainability, University of Minho, Campus de Gualtar, 4710-057, Braga, Portugal
^3^Centro de Química Estrutural, Institute of Molecular Sciences and Departamento de Química e Bioquímica, Faculdade de Ciências, Universidade de Lisboa, Campo Grande, 1749-016 Lisboa, Portugal
^4^Institute for Research and Innovation in Health (i3S), University of Porto, 4200-135 Porto, Portugal
^5^Institute of Molecular Pathology and Immunology of the University of Porto, 4200-135 Porto, Portugal
^6^BioISI - Instituto de Biosistemas e Ciências Integrativas and Departamento de Química e Bioquímica, Faculdade de Ciências, Universidade de Lisboa, 1749-016, Lisboa, Portugal
^7^Department of Pathological, Cytological and Thanatological Anatomy, School of Health, Polytechnic Institute of Porto, 4200-072 Porto, Portugal
^8^Life and Health Sciences Research Institute (ICVS), School of Medicine, University of Minho, 4710-057 Braga, Portugal
^9^ICVS/3B’s-PT Government Associate Laboratory, 4710-057 Braga, Portugal
^10^ICVS/3B’s-PT Government Associate Laboratory, 4806-909 Guimarães, Portugal
^11^Center for Interdisciplinary Research Egas Moniz (CiiEM), Quinta da Granja, Monte da Caparica, 2829-511 Caparica, Portugal.
^12^Instituto de Anatomia Patológica, Faculdade de Medicina da Universidade de Lisboa, Av. Prof. Egas Moniz, 1649-028 Lisboa, Portugal
^13^Institute of Biomedical Sciences of Abel Salazar, University of Porto, 4050-313 Porto, Portugal
^#^These authors share senior authorship

### Slide 2
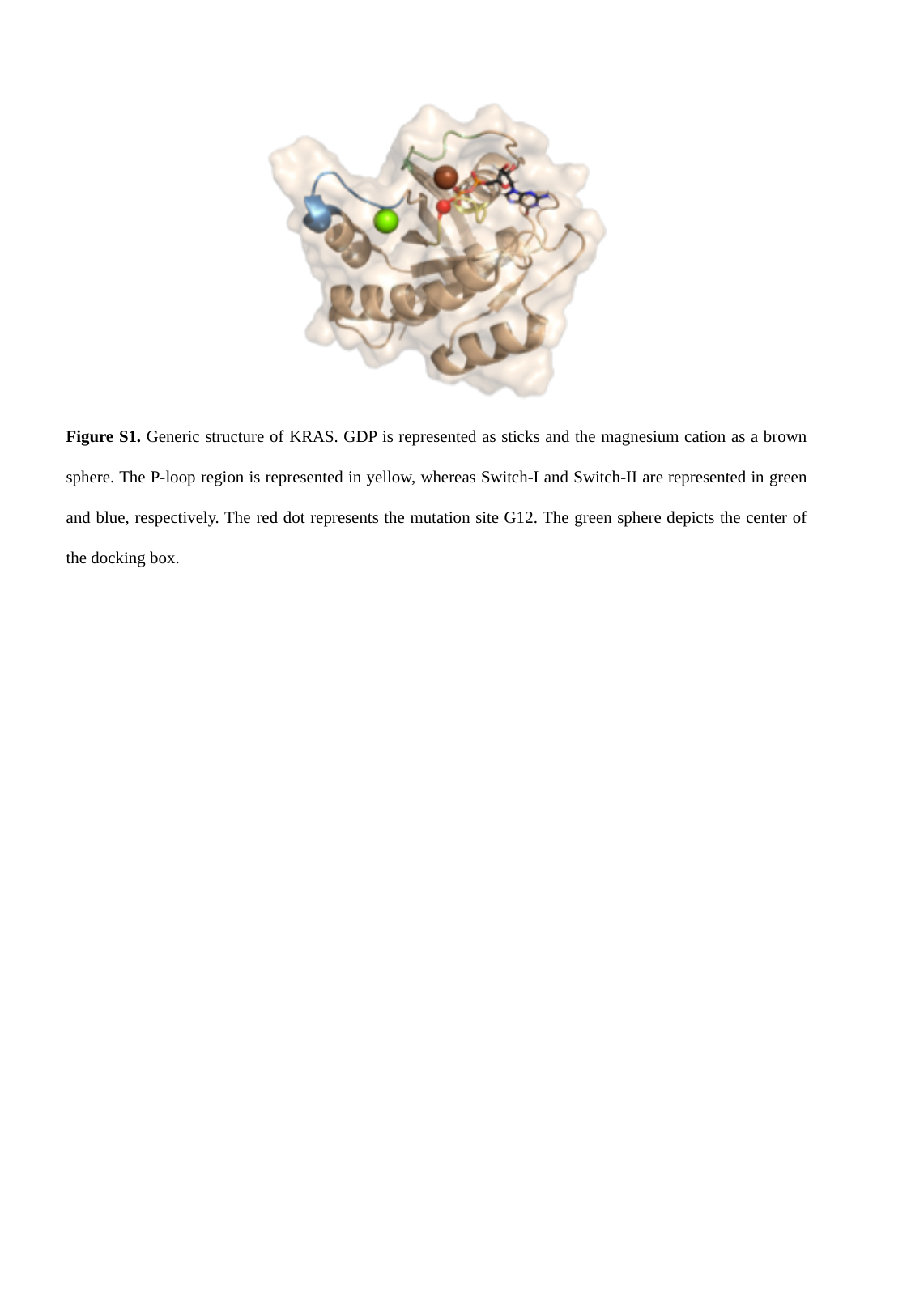

Figure S1. Generic structure of KRAS. GDP is represented as sticks and the magnesium cation as a brown sphere. The P-loop region is represented in yellow, whereas Switch-I and Switch-II are represented in green and blue, respectively. The red dot represents the mutation site G12. The green sphere depicts the center of the docking box.

### Slide 3
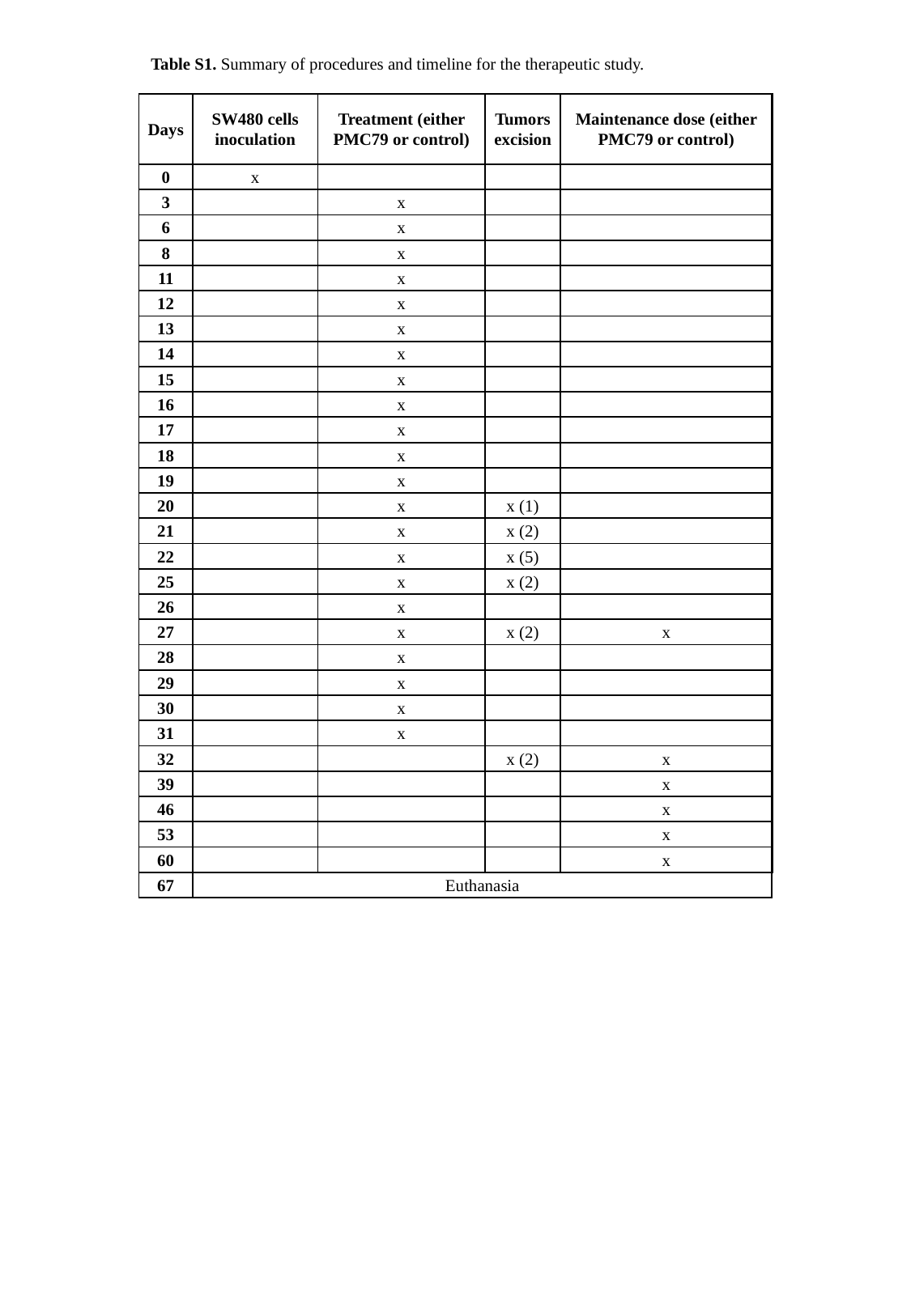

Table S1. Summary of procedures and timeline for the therapeutic study.
| Days | SW480 cells inoculation | Treatment (either PMC79 or control) | Tumors excision | Maintenance dose (either PMC79 or control) |
| --- | --- | --- | --- | --- |
| 0 | x | | | |
| 3 | | x | | |
| 6 | | x | | |
| 8 | | x | | |
| 11 | | x | | |
| 12 | | x | | |
| 13 | | x | | |
| 14 | | x | | |
| 15 | | x | | |
| 16 | | x | | |
| 17 | | x | | |
| 18 | | x | | |
| 19 | | x | | |
| 20 | | x | x (1) | |
| 21 | | x | x (2) | |
| 22 | | x | x (5) | |
| 25 | | x | x (2) | |
| 26 | | x | | |
| 27 | | x | x (2) | x |
| 28 | | x | | |
| 29 | | x | | |
| 30 | | x | | |
| 31 | | x | | |
| 32 | | | x (2) | x |
| 39 | | | | x |
| 46 | | | | x |
| 53 | | | | x |
| 60 | | | | x |
| 67 | Euthanasia | | | |

### Slide 4
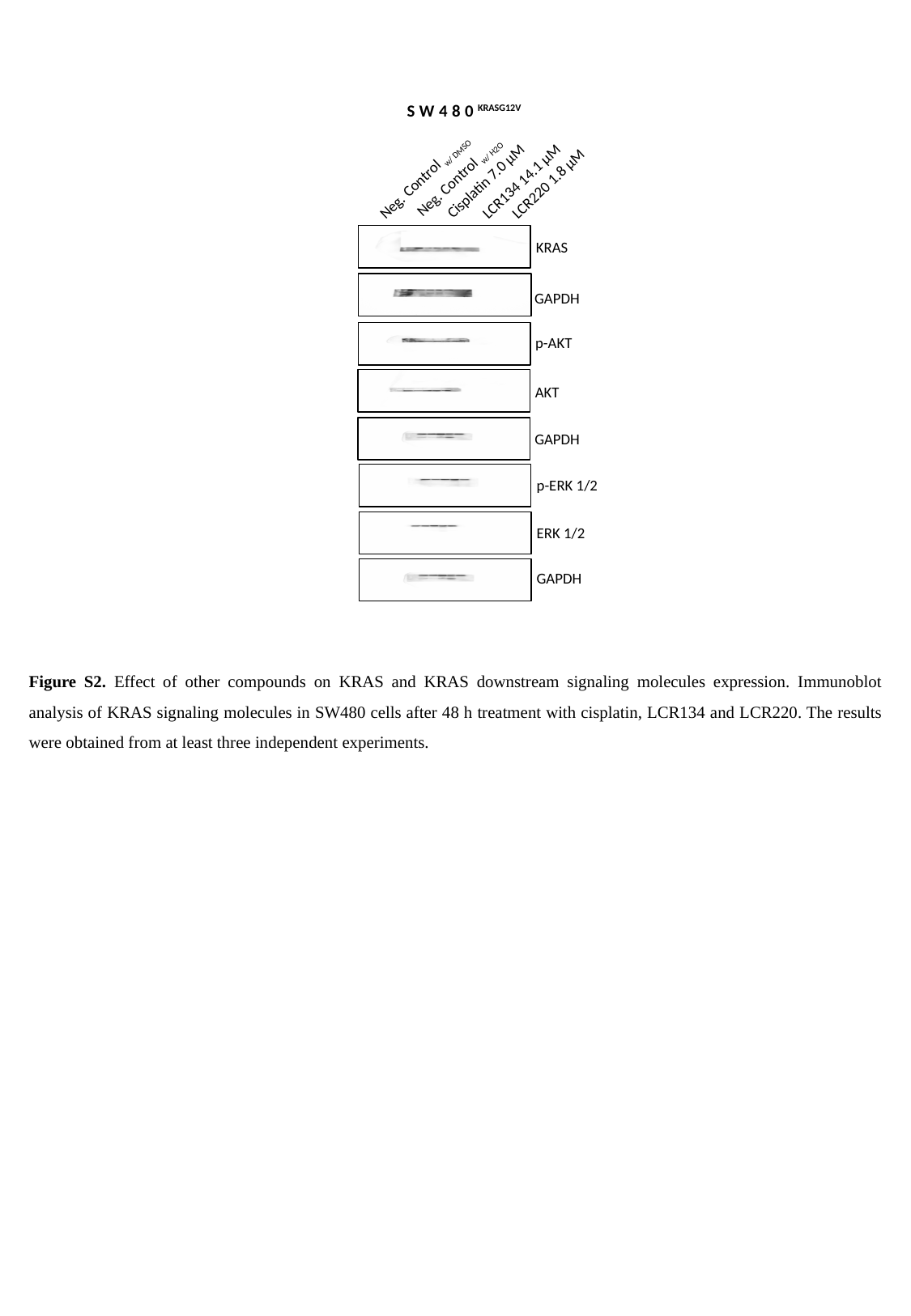

SW480^KRASG12V^
Neg. Control _w/ H2O_
Neg. Control _w/ DMSO_
LCR220 1.8 µM
LCR134 14.1 µM
Cisplatin 7.0 µM
KRAS
GAPDH
p-AKT
AKT
GAPDH
p-ERK 1/2
ERK 1/2
GAPDH
Figure S2. Effect of other compounds on KRAS and KRAS downstream signaling molecules expression. Immunoblot analysis of KRAS signaling molecules in SW480 cells after 48 h treatment with cisplatin, LCR134 and LCR220. The results were obtained from at least three independent experiments.

### Slide 5
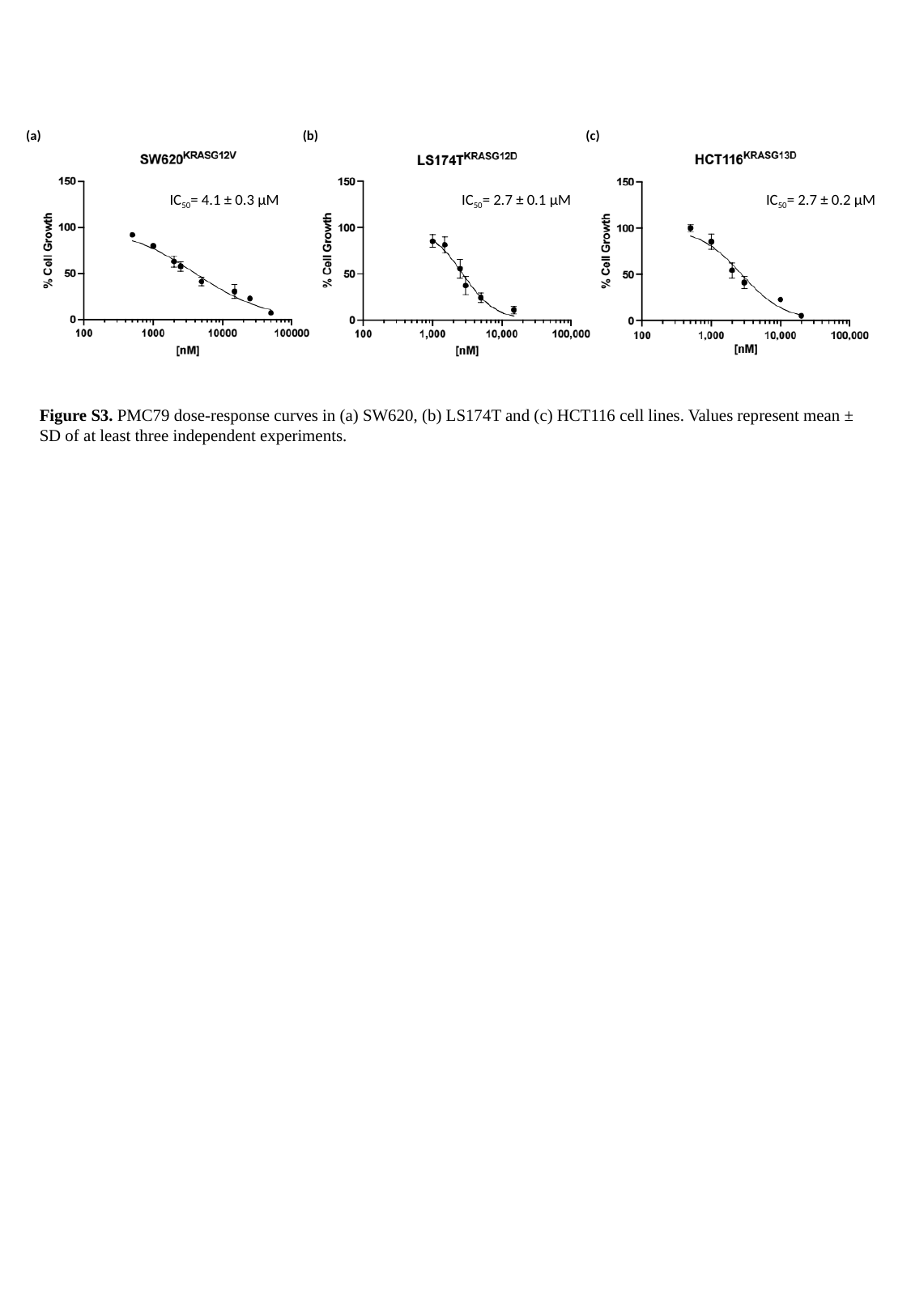

(a)
(b)
(c)
IC_50_= 4.1 ± 0.3 μM
IC_50_= 2.7 ± 0.1 μM
IC_50_= 2.7 ± 0.2 μM
Figure S3. PMC79 dose-response curves in (a) SW620, (b) LS174T and (c) HCT116 cell lines. Values represent mean ± SD of at least three independent experiments.

### Slide 6
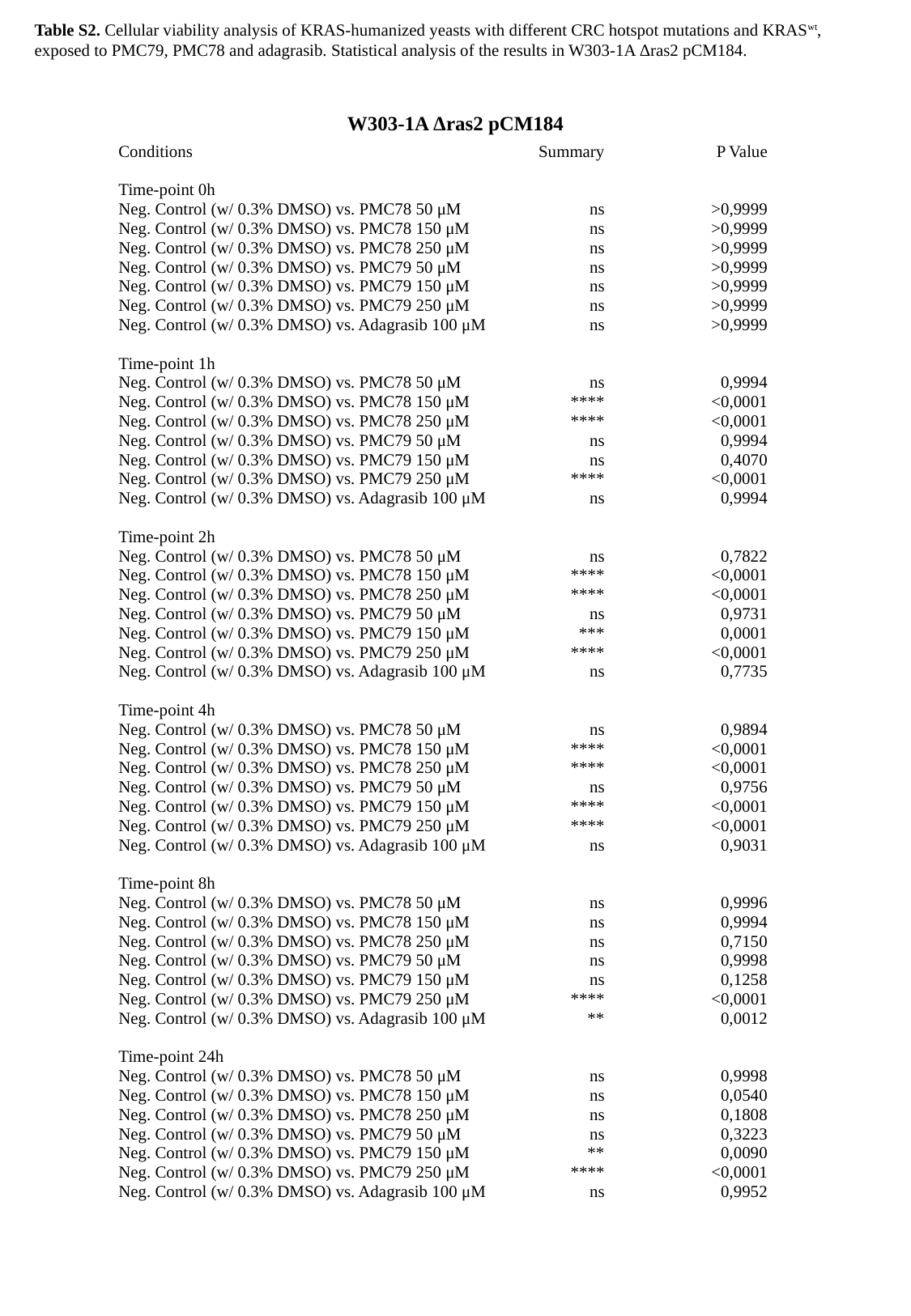

Table S2. Cellular viability analysis of KRAS-humanized yeasts with different CRC hotspot mutations and KRAS^wt^, exposed to PMC79, PMC78 and adagrasib. Statistical analysis of the results in W303-1A Δras2 pCM184.
W303-1A Δras2 pCM184
| Conditions | Summary | P Value |
| --- | --- | --- |
| Time-point 0h | | |
| Neg. Control (w/ 0.3% DMSO) vs. PMC78 50 μM | ns | >0,9999 |
| Neg. Control (w/ 0.3% DMSO) vs. PMC78 150 μM | ns | >0,9999 |
| Neg. Control (w/ 0.3% DMSO) vs. PMC78 250 μM | ns | >0,9999 |
| Neg. Control (w/ 0.3% DMSO) vs. PMC79 50 μM | ns | >0,9999 |
| Neg. Control (w/ 0.3% DMSO) vs. PMC79 150 μM | ns | >0,9999 |
| Neg. Control (w/ 0.3% DMSO) vs. PMC79 250 μM | ns | >0,9999 |
| Neg. Control (w/ 0.3% DMSO) vs. Adagrasib 100 μM | ns | >0,9999 |
| Time-point 1h | | |
| Neg. Control (w/ 0.3% DMSO) vs. PMC78 50 μM | ns | 0,9994 |
| Neg. Control (w/ 0.3% DMSO) vs. PMC78 150 μM | \*\*\*\* | <0,0001 |
| Neg. Control (w/ 0.3% DMSO) vs. PMC78 250 μM | \*\*\*\* | <0,0001 |
| Neg. Control (w/ 0.3% DMSO) vs. PMC79 50 μM | ns | 0,9994 |
| Neg. Control (w/ 0.3% DMSO) vs. PMC79 150 μM | ns | 0,4070 |
| Neg. Control (w/ 0.3% DMSO) vs. PMC79 250 μM | \*\*\*\* | <0,0001 |
| Neg. Control (w/ 0.3% DMSO) vs. Adagrasib 100 μM | ns | 0,9994 |
| Time-point 2h | | |
| Neg. Control (w/ 0.3% DMSO) vs. PMC78 50 μM | ns | 0,7822 |
| Neg. Control (w/ 0.3% DMSO) vs. PMC78 150 μM | \*\*\*\* | <0,0001 |
| Neg. Control (w/ 0.3% DMSO) vs. PMC78 250 μM | \*\*\*\* | <0,0001 |
| Neg. Control (w/ 0.3% DMSO) vs. PMC79 50 μM | ns | 0,9731 |
| Neg. Control (w/ 0.3% DMSO) vs. PMC79 150 μM | \*\*\* | 0,0001 |
| Neg. Control (w/ 0.3% DMSO) vs. PMC79 250 μM | \*\*\*\* | <0,0001 |
| Neg. Control (w/ 0.3% DMSO) vs. Adagrasib 100 μM | ns | 0,7735 |
| Time-point 4h | | |
| Neg. Control (w/ 0.3% DMSO) vs. PMC78 50 μM | ns | 0,9894 |
| Neg. Control (w/ 0.3% DMSO) vs. PMC78 150 μM | \*\*\*\* | <0,0001 |
| Neg. Control (w/ 0.3% DMSO) vs. PMC78 250 μM | \*\*\*\* | <0,0001 |
| Neg. Control (w/ 0.3% DMSO) vs. PMC79 50 μM | ns | 0,9756 |
| Neg. Control (w/ 0.3% DMSO) vs. PMC79 150 μM | \*\*\*\* | <0,0001 |
| Neg. Control (w/ 0.3% DMSO) vs. PMC79 250 μM | \*\*\*\* | <0,0001 |
| Neg. Control (w/ 0.3% DMSO) vs. Adagrasib 100 μM | ns | 0,9031 |
| Time-point 8h | | |
| Neg. Control (w/ 0.3% DMSO) vs. PMC78 50 μM | ns | 0,9996 |
| Neg. Control (w/ 0.3% DMSO) vs. PMC78 150 μM | ns | 0,9994 |
| Neg. Control (w/ 0.3% DMSO) vs. PMC78 250 μM | ns | 0,7150 |
| Neg. Control (w/ 0.3% DMSO) vs. PMC79 50 μM | ns | 0,9998 |
| Neg. Control (w/ 0.3% DMSO) vs. PMC79 150 μM | ns | 0,1258 |
| Neg. Control (w/ 0.3% DMSO) vs. PMC79 250 μM | \*\*\*\* | <0,0001 |
| Neg. Control (w/ 0.3% DMSO) vs. Adagrasib 100 μM | \*\* | 0,0012 |
| Time-point 24h | | |
| Neg. Control (w/ 0.3% DMSO) vs. PMC78 50 μM | ns | 0,9998 |
| Neg. Control (w/ 0.3% DMSO) vs. PMC78 150 μM | ns | 0,0540 |
| Neg. Control (w/ 0.3% DMSO) vs. PMC78 250 μM | ns | 0,1808 |
| Neg. Control (w/ 0.3% DMSO) vs. PMC79 50 μM | ns | 0,3223 |
| Neg. Control (w/ 0.3% DMSO) vs. PMC79 150 μM | \*\* | 0,0090 |
| Neg. Control (w/ 0.3% DMSO) vs. PMC79 250 μM | \*\*\*\* | <0,0001 |
| Neg. Control (w/ 0.3% DMSO) vs. Adagrasib 100 μM | ns | 0,9952 |

### Slide 7
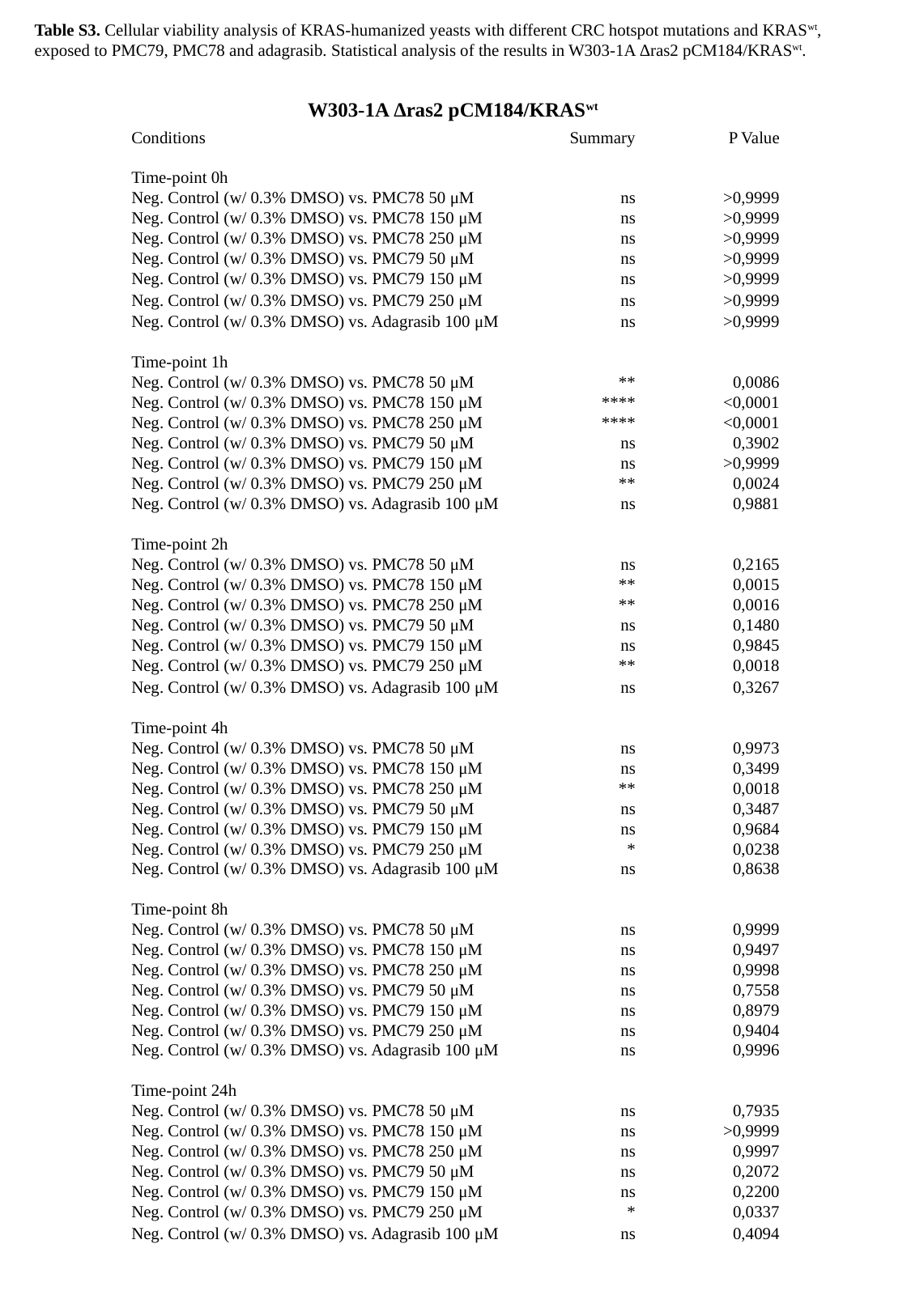

Table S3. Cellular viability analysis of KRAS-humanized yeasts with different CRC hotspot mutations and KRAS^wt^, exposed to PMC79, PMC78 and adagrasib. Statistical analysis of the results in W303-1A Δras2 pCM184/KRAS^wt^.
W303-1A Δras2 pCM184/KRAS^wt^
| Conditions | Summary | P Value |
| --- | --- | --- |
| Time-point 0h | | |
| Neg. Control (w/ 0.3% DMSO) vs. PMC78 50 μM | ns | >0,9999 |
| Neg. Control (w/ 0.3% DMSO) vs. PMC78 150 μM | ns | >0,9999 |
| Neg. Control (w/ 0.3% DMSO) vs. PMC78 250 μM | ns | >0,9999 |
| Neg. Control (w/ 0.3% DMSO) vs. PMC79 50 μM | ns | >0,9999 |
| Neg. Control (w/ 0.3% DMSO) vs. PMC79 150 μM | ns | >0,9999 |
| Neg. Control (w/ 0.3% DMSO) vs. PMC79 250 μM | ns | >0,9999 |
| Neg. Control (w/ 0.3% DMSO) vs. Adagrasib 100 μM | ns | >0,9999 |
| Time-point 1h | | |
| Neg. Control (w/ 0.3% DMSO) vs. PMC78 50 μM | \*\* | 0,0086 |
| Neg. Control (w/ 0.3% DMSO) vs. PMC78 150 μM | \*\*\*\* | <0,0001 |
| Neg. Control (w/ 0.3% DMSO) vs. PMC78 250 μM | \*\*\*\* | <0,0001 |
| Neg. Control (w/ 0.3% DMSO) vs. PMC79 50 μM | ns | 0,3902 |
| Neg. Control (w/ 0.3% DMSO) vs. PMC79 150 μM | ns | >0,9999 |
| Neg. Control (w/ 0.3% DMSO) vs. PMC79 250 μM | \*\* | 0,0024 |
| Neg. Control (w/ 0.3% DMSO) vs. Adagrasib 100 μM | ns | 0,9881 |
| Time-point 2h | | |
| Neg. Control (w/ 0.3% DMSO) vs. PMC78 50 μM | ns | 0,2165 |
| Neg. Control (w/ 0.3% DMSO) vs. PMC78 150 μM | \*\* | 0,0015 |
| Neg. Control (w/ 0.3% DMSO) vs. PMC78 250 μM | \*\* | 0,0016 |
| Neg. Control (w/ 0.3% DMSO) vs. PMC79 50 μM | ns | 0,1480 |
| Neg. Control (w/ 0.3% DMSO) vs. PMC79 150 μM | ns | 0,9845 |
| Neg. Control (w/ 0.3% DMSO) vs. PMC79 250 μM | \*\* | 0,0018 |
| Neg. Control (w/ 0.3% DMSO) vs. Adagrasib 100 μM | ns | 0,3267 |
| Time-point 4h | | |
| Neg. Control (w/ 0.3% DMSO) vs. PMC78 50 μM | ns | 0,9973 |
| Neg. Control (w/ 0.3% DMSO) vs. PMC78 150 μM | ns | 0,3499 |
| Neg. Control (w/ 0.3% DMSO) vs. PMC78 250 μM | \*\* | 0,0018 |
| Neg. Control (w/ 0.3% DMSO) vs. PMC79 50 μM | ns | 0,3487 |
| Neg. Control (w/ 0.3% DMSO) vs. PMC79 150 μM | ns | 0,9684 |
| Neg. Control (w/ 0.3% DMSO) vs. PMC79 250 μM | \* | 0,0238 |
| Neg. Control (w/ 0.3% DMSO) vs. Adagrasib 100 μM | ns | 0,8638 |
| Time-point 8h | | |
| Neg. Control (w/ 0.3% DMSO) vs. PMC78 50 μM | ns | 0,9999 |
| Neg. Control (w/ 0.3% DMSO) vs. PMC78 150 μM | ns | 0,9497 |
| Neg. Control (w/ 0.3% DMSO) vs. PMC78 250 μM | ns | 0,9998 |
| Neg. Control (w/ 0.3% DMSO) vs. PMC79 50 μM | ns | 0,7558 |
| Neg. Control (w/ 0.3% DMSO) vs. PMC79 150 μM | ns | 0,8979 |
| Neg. Control (w/ 0.3% DMSO) vs. PMC79 250 μM | ns | 0,9404 |
| Neg. Control (w/ 0.3% DMSO) vs. Adagrasib 100 μM | ns | 0,9996 |
| Time-point 24h | | |
| Neg. Control (w/ 0.3% DMSO) vs. PMC78 50 μM | ns | 0,7935 |
| Neg. Control (w/ 0.3% DMSO) vs. PMC78 150 μM | ns | >0,9999 |
| Neg. Control (w/ 0.3% DMSO) vs. PMC78 250 μM | ns | 0,9997 |
| Neg. Control (w/ 0.3% DMSO) vs. PMC79 50 μM | ns | 0,2072 |
| Neg. Control (w/ 0.3% DMSO) vs. PMC79 150 μM | ns | 0,2200 |
| Neg. Control (w/ 0.3% DMSO) vs. PMC79 250 μM | \* | 0,0337 |
| Neg. Control (w/ 0.3% DMSO) vs. Adagrasib 100 μM | ns | 0,4094 |

### Slide 8
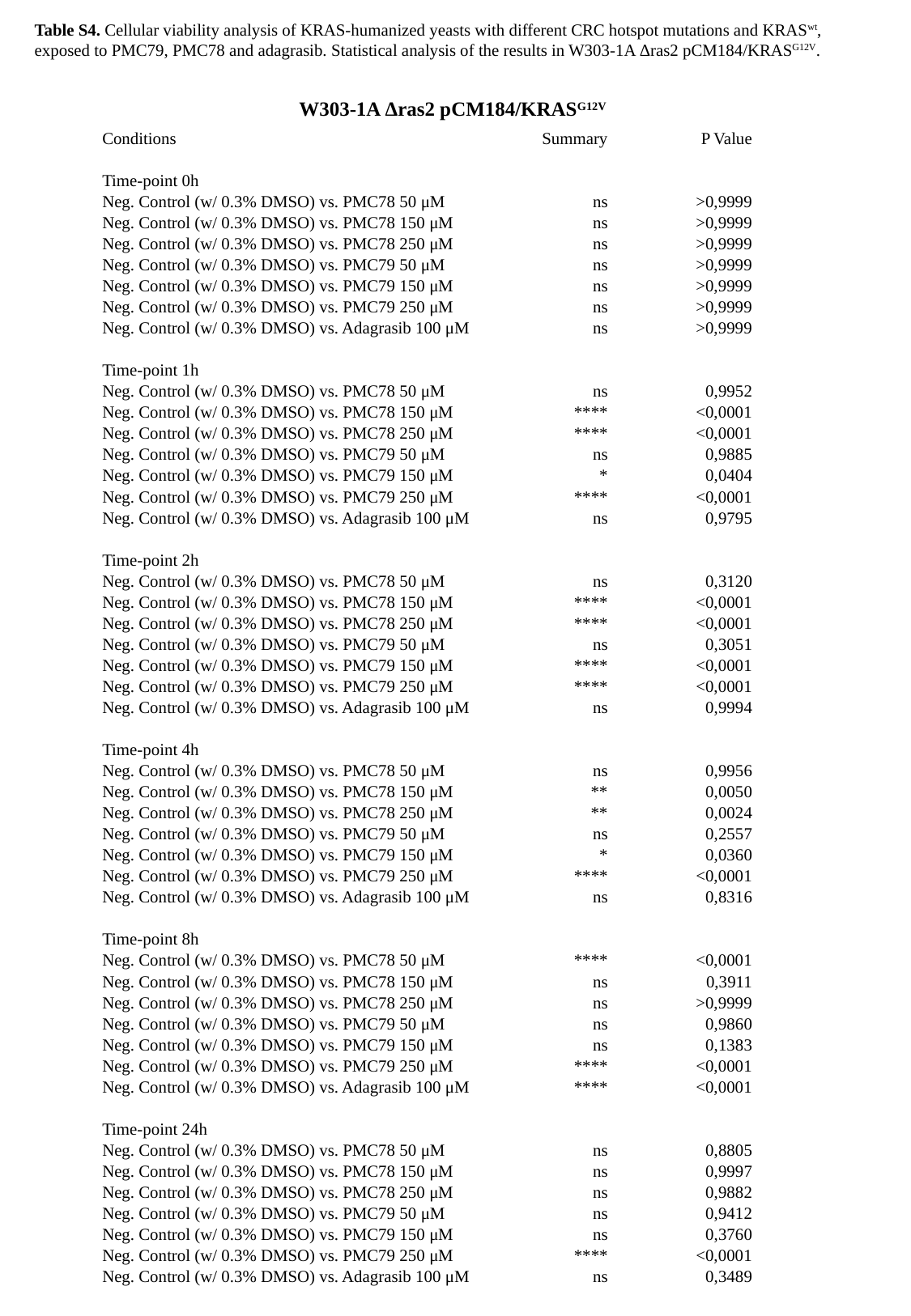

Table S4. Cellular viability analysis of KRAS-humanized yeasts with different CRC hotspot mutations and KRAS^wt^, exposed to PMC79, PMC78 and adagrasib. Statistical analysis of the results in W303-1A Δras2 pCM184/KRAS^G12V^.
W303-1A Δras2 pCM184/KRAS^G12V^
| Conditions | Summary | P Value |
| --- | --- | --- |
| Time-point 0h | | |
| Neg. Control (w/ 0.3% DMSO) vs. PMC78 50 μM | ns | >0,9999 |
| Neg. Control (w/ 0.3% DMSO) vs. PMC78 150 μM | ns | >0,9999 |
| Neg. Control (w/ 0.3% DMSO) vs. PMC78 250 μM | ns | >0,9999 |
| Neg. Control (w/ 0.3% DMSO) vs. PMC79 50 μM | ns | >0,9999 |
| Neg. Control (w/ 0.3% DMSO) vs. PMC79 150 μM | ns | >0,9999 |
| Neg. Control (w/ 0.3% DMSO) vs. PMC79 250 μM | ns | >0,9999 |
| Neg. Control (w/ 0.3% DMSO) vs. Adagrasib 100 μM | ns | >0,9999 |
| Time-point 1h | | |
| Neg. Control (w/ 0.3% DMSO) vs. PMC78 50 μM | ns | 0,9952 |
| Neg. Control (w/ 0.3% DMSO) vs. PMC78 150 μM | \*\*\*\* | <0,0001 |
| Neg. Control (w/ 0.3% DMSO) vs. PMC78 250 μM | \*\*\*\* | <0,0001 |
| Neg. Control (w/ 0.3% DMSO) vs. PMC79 50 μM | ns | 0,9885 |
| Neg. Control (w/ 0.3% DMSO) vs. PMC79 150 μM | \* | 0,0404 |
| Neg. Control (w/ 0.3% DMSO) vs. PMC79 250 μM | \*\*\*\* | <0,0001 |
| Neg. Control (w/ 0.3% DMSO) vs. Adagrasib 100 μM | ns | 0,9795 |
| Time-point 2h | | |
| Neg. Control (w/ 0.3% DMSO) vs. PMC78 50 μM | ns | 0,3120 |
| Neg. Control (w/ 0.3% DMSO) vs. PMC78 150 μM | \*\*\*\* | <0,0001 |
| Neg. Control (w/ 0.3% DMSO) vs. PMC78 250 μM | \*\*\*\* | <0,0001 |
| Neg. Control (w/ 0.3% DMSO) vs. PMC79 50 μM | ns | 0,3051 |
| Neg. Control (w/ 0.3% DMSO) vs. PMC79 150 μM | \*\*\*\* | <0,0001 |
| Neg. Control (w/ 0.3% DMSO) vs. PMC79 250 μM | \*\*\*\* | <0,0001 |
| Neg. Control (w/ 0.3% DMSO) vs. Adagrasib 100 μM | ns | 0,9994 |
| Time-point 4h | | |
| Neg. Control (w/ 0.3% DMSO) vs. PMC78 50 μM | ns | 0,9956 |
| Neg. Control (w/ 0.3% DMSO) vs. PMC78 150 μM | \*\* | 0,0050 |
| Neg. Control (w/ 0.3% DMSO) vs. PMC78 250 μM | \*\* | 0,0024 |
| Neg. Control (w/ 0.3% DMSO) vs. PMC79 50 μM | ns | 0,2557 |
| Neg. Control (w/ 0.3% DMSO) vs. PMC79 150 μM | \* | 0,0360 |
| Neg. Control (w/ 0.3% DMSO) vs. PMC79 250 μM | \*\*\*\* | <0,0001 |
| Neg. Control (w/ 0.3% DMSO) vs. Adagrasib 100 μM | ns | 0,8316 |
| Time-point 8h | | |
| Neg. Control (w/ 0.3% DMSO) vs. PMC78 50 μM | \*\*\*\* | <0,0001 |
| Neg. Control (w/ 0.3% DMSO) vs. PMC78 150 μM | ns | 0,3911 |
| Neg. Control (w/ 0.3% DMSO) vs. PMC78 250 μM | ns | >0,9999 |
| Neg. Control (w/ 0.3% DMSO) vs. PMC79 50 μM | ns | 0,9860 |
| Neg. Control (w/ 0.3% DMSO) vs. PMC79 150 μM | ns | 0,1383 |
| Neg. Control (w/ 0.3% DMSO) vs. PMC79 250 μM | \*\*\*\* | <0,0001 |
| Neg. Control (w/ 0.3% DMSO) vs. Adagrasib 100 μM | \*\*\*\* | <0,0001 |
| Time-point 24h | | |
| Neg. Control (w/ 0.3% DMSO) vs. PMC78 50 μM | ns | 0,8805 |
| Neg. Control (w/ 0.3% DMSO) vs. PMC78 150 μM | ns | 0,9997 |
| Neg. Control (w/ 0.3% DMSO) vs. PMC78 250 μM | ns | 0,9882 |
| Neg. Control (w/ 0.3% DMSO) vs. PMC79 50 μM | ns | 0,9412 |
| Neg. Control (w/ 0.3% DMSO) vs. PMC79 150 μM | ns | 0,3760 |
| Neg. Control (w/ 0.3% DMSO) vs. PMC79 250 μM | \*\*\*\* | <0,0001 |
| Neg. Control (w/ 0.3% DMSO) vs. Adagrasib 100 μM | ns | 0,3489 |

### Slide 9
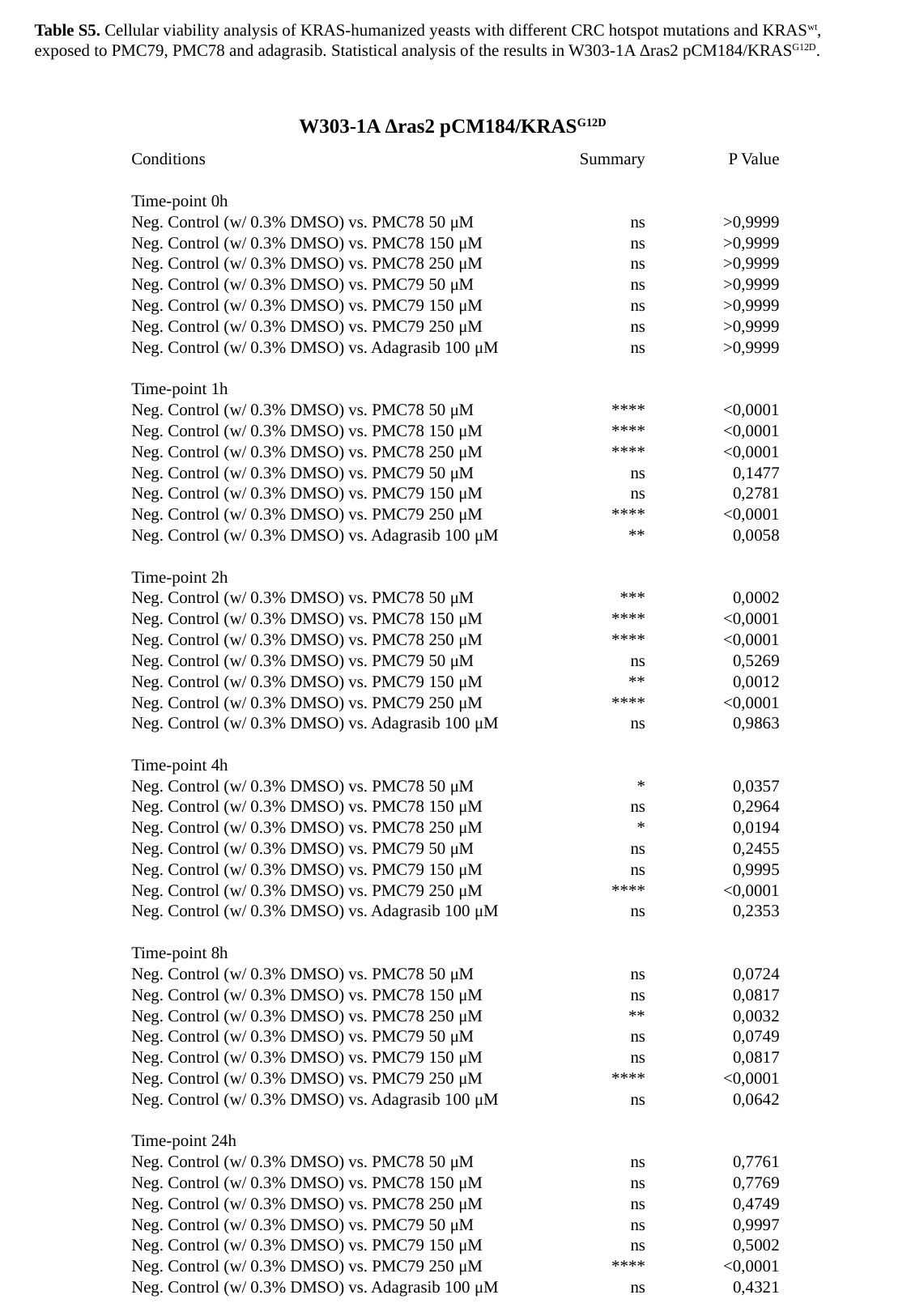

Table S5. Cellular viability analysis of KRAS-humanized yeasts with different CRC hotspot mutations and KRAS^wt^, exposed to PMC79, PMC78 and adagrasib. Statistical analysis of the results in W303-1A Δras2 pCM184/KRAS^G12D^.
W303-1A Δras2 pCM184/KRAS^G12D^
| Conditions | Summary | P Value |
| --- | --- | --- |
| Time-point 0h | | |
| Neg. Control (w/ 0.3% DMSO) vs. PMC78 50 μM | ns | >0,9999 |
| Neg. Control (w/ 0.3% DMSO) vs. PMC78 150 μM | ns | >0,9999 |
| Neg. Control (w/ 0.3% DMSO) vs. PMC78 250 μM | ns | >0,9999 |
| Neg. Control (w/ 0.3% DMSO) vs. PMC79 50 μM | ns | >0,9999 |
| Neg. Control (w/ 0.3% DMSO) vs. PMC79 150 μM | ns | >0,9999 |
| Neg. Control (w/ 0.3% DMSO) vs. PMC79 250 μM | ns | >0,9999 |
| Neg. Control (w/ 0.3% DMSO) vs. Adagrasib 100 μM | ns | >0,9999 |
| Time-point 1h | | |
| Neg. Control (w/ 0.3% DMSO) vs. PMC78 50 μM | \*\*\*\* | <0,0001 |
| Neg. Control (w/ 0.3% DMSO) vs. PMC78 150 μM | \*\*\*\* | <0,0001 |
| Neg. Control (w/ 0.3% DMSO) vs. PMC78 250 μM | \*\*\*\* | <0,0001 |
| Neg. Control (w/ 0.3% DMSO) vs. PMC79 50 μM | ns | 0,1477 |
| Neg. Control (w/ 0.3% DMSO) vs. PMC79 150 μM | ns | 0,2781 |
| Neg. Control (w/ 0.3% DMSO) vs. PMC79 250 μM | \*\*\*\* | <0,0001 |
| Neg. Control (w/ 0.3% DMSO) vs. Adagrasib 100 μM | \*\* | 0,0058 |
| Time-point 2h | | |
| Neg. Control (w/ 0.3% DMSO) vs. PMC78 50 μM | \*\*\* | 0,0002 |
| Neg. Control (w/ 0.3% DMSO) vs. PMC78 150 μM | \*\*\*\* | <0,0001 |
| Neg. Control (w/ 0.3% DMSO) vs. PMC78 250 μM | \*\*\*\* | <0,0001 |
| Neg. Control (w/ 0.3% DMSO) vs. PMC79 50 μM | ns | 0,5269 |
| Neg. Control (w/ 0.3% DMSO) vs. PMC79 150 μM | \*\* | 0,0012 |
| Neg. Control (w/ 0.3% DMSO) vs. PMC79 250 μM | \*\*\*\* | <0,0001 |
| Neg. Control (w/ 0.3% DMSO) vs. Adagrasib 100 μM | ns | 0,9863 |
| Time-point 4h | | |
| Neg. Control (w/ 0.3% DMSO) vs. PMC78 50 μM | \* | 0,0357 |
| Neg. Control (w/ 0.3% DMSO) vs. PMC78 150 μM | ns | 0,2964 |
| Neg. Control (w/ 0.3% DMSO) vs. PMC78 250 μM | \* | 0,0194 |
| Neg. Control (w/ 0.3% DMSO) vs. PMC79 50 μM | ns | 0,2455 |
| Neg. Control (w/ 0.3% DMSO) vs. PMC79 150 μM | ns | 0,9995 |
| Neg. Control (w/ 0.3% DMSO) vs. PMC79 250 μM | \*\*\*\* | <0,0001 |
| Neg. Control (w/ 0.3% DMSO) vs. Adagrasib 100 μM | ns | 0,2353 |
| Time-point 8h | | |
| Neg. Control (w/ 0.3% DMSO) vs. PMC78 50 μM | ns | 0,0724 |
| Neg. Control (w/ 0.3% DMSO) vs. PMC78 150 μM | ns | 0,0817 |
| Neg. Control (w/ 0.3% DMSO) vs. PMC78 250 μM | \*\* | 0,0032 |
| Neg. Control (w/ 0.3% DMSO) vs. PMC79 50 μM | ns | 0,0749 |
| Neg. Control (w/ 0.3% DMSO) vs. PMC79 150 μM | ns | 0,0817 |
| Neg. Control (w/ 0.3% DMSO) vs. PMC79 250 μM | \*\*\*\* | <0,0001 |
| Neg. Control (w/ 0.3% DMSO) vs. Adagrasib 100 μM | ns | 0,0642 |
| Time-point 24h | | |
| Neg. Control (w/ 0.3% DMSO) vs. PMC78 50 μM | ns | 0,7761 |
| Neg. Control (w/ 0.3% DMSO) vs. PMC78 150 μM | ns | 0,7769 |
| Neg. Control (w/ 0.3% DMSO) vs. PMC78 250 μM | ns | 0,4749 |
| Neg. Control (w/ 0.3% DMSO) vs. PMC79 50 μM | ns | 0,9997 |
| Neg. Control (w/ 0.3% DMSO) vs. PMC79 150 μM | ns | 0,5002 |
| Neg. Control (w/ 0.3% DMSO) vs. PMC79 250 μM | \*\*\*\* | <0,0001 |
| Neg. Control (w/ 0.3% DMSO) vs. Adagrasib 100 μM | ns | 0,4321 |

### Slide 10
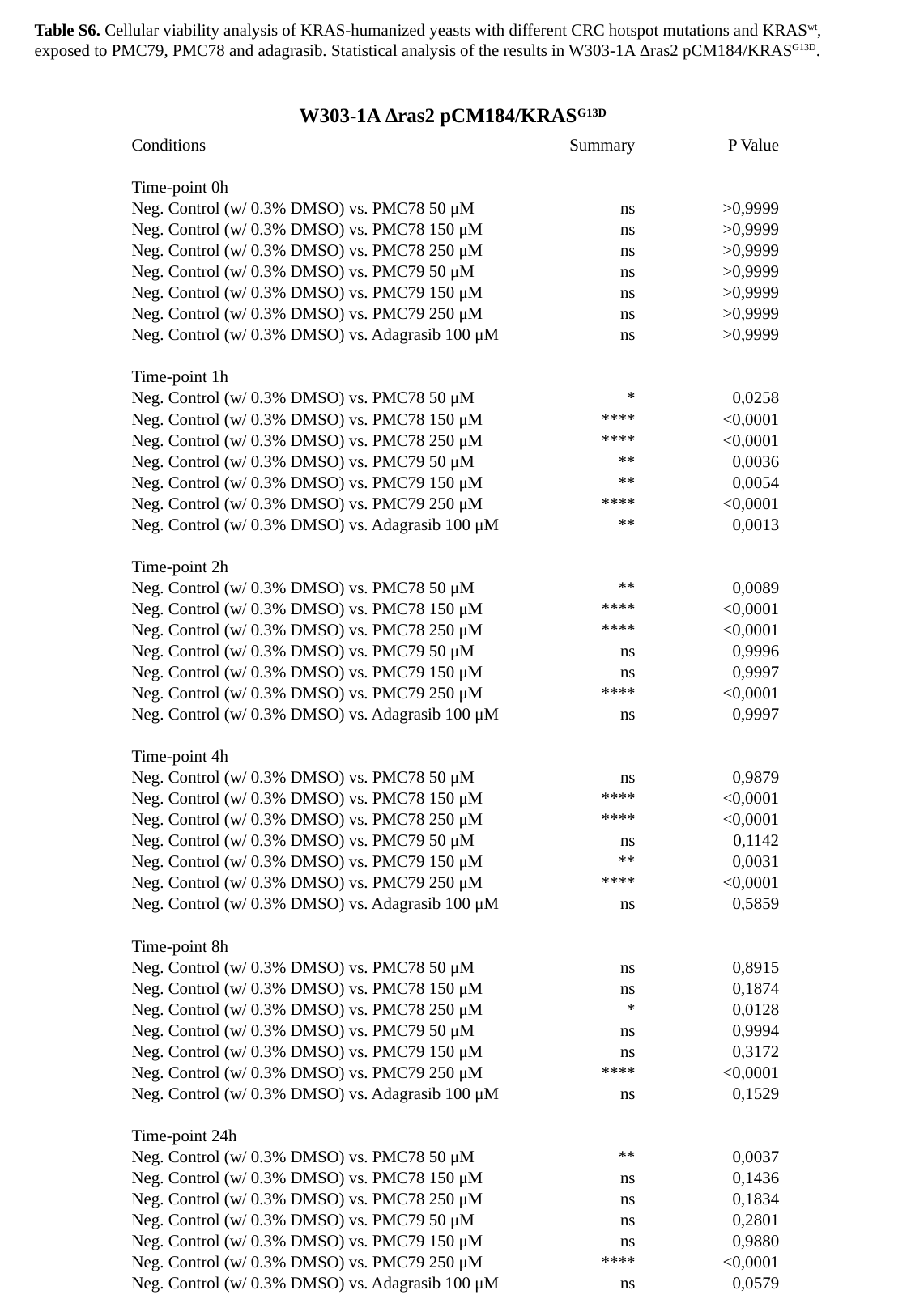

Table S6. Cellular viability analysis of KRAS-humanized yeasts with different CRC hotspot mutations and KRAS^wt^, exposed to PMC79, PMC78 and adagrasib. Statistical analysis of the results in W303-1A Δras2 pCM184/KRAS^G13D^.
W303-1A Δras2 pCM184/KRAS^G13D^
| Conditions | Summary | P Value |
| --- | --- | --- |
| Time-point 0h | | |
| Neg. Control (w/ 0.3% DMSO) vs. PMC78 50 μM | ns | >0,9999 |
| Neg. Control (w/ 0.3% DMSO) vs. PMC78 150 μM | ns | >0,9999 |
| Neg. Control (w/ 0.3% DMSO) vs. PMC78 250 μM | ns | >0,9999 |
| Neg. Control (w/ 0.3% DMSO) vs. PMC79 50 μM | ns | >0,9999 |
| Neg. Control (w/ 0.3% DMSO) vs. PMC79 150 μM | ns | >0,9999 |
| Neg. Control (w/ 0.3% DMSO) vs. PMC79 250 μM | ns | >0,9999 |
| Neg. Control (w/ 0.3% DMSO) vs. Adagrasib 100 μM | ns | >0,9999 |
| Time-point 1h | | |
| Neg. Control (w/ 0.3% DMSO) vs. PMC78 50 μM | \* | 0,0258 |
| Neg. Control (w/ 0.3% DMSO) vs. PMC78 150 μM | \*\*\*\* | <0,0001 |
| Neg. Control (w/ 0.3% DMSO) vs. PMC78 250 μM | \*\*\*\* | <0,0001 |
| Neg. Control (w/ 0.3% DMSO) vs. PMC79 50 μM | \*\* | 0,0036 |
| Neg. Control (w/ 0.3% DMSO) vs. PMC79 150 μM | \*\* | 0,0054 |
| Neg. Control (w/ 0.3% DMSO) vs. PMC79 250 μM | \*\*\*\* | <0,0001 |
| Neg. Control (w/ 0.3% DMSO) vs. Adagrasib 100 μM | \*\* | 0,0013 |
| Time-point 2h | | |
| Neg. Control (w/ 0.3% DMSO) vs. PMC78 50 μM | \*\* | 0,0089 |
| Neg. Control (w/ 0.3% DMSO) vs. PMC78 150 μM | \*\*\*\* | <0,0001 |
| Neg. Control (w/ 0.3% DMSO) vs. PMC78 250 μM | \*\*\*\* | <0,0001 |
| Neg. Control (w/ 0.3% DMSO) vs. PMC79 50 μM | ns | 0,9996 |
| Neg. Control (w/ 0.3% DMSO) vs. PMC79 150 μM | ns | 0,9997 |
| Neg. Control (w/ 0.3% DMSO) vs. PMC79 250 μM | \*\*\*\* | <0,0001 |
| Neg. Control (w/ 0.3% DMSO) vs. Adagrasib 100 μM | ns | 0,9997 |
| Time-point 4h | | |
| Neg. Control (w/ 0.3% DMSO) vs. PMC78 50 μM | ns | 0,9879 |
| Neg. Control (w/ 0.3% DMSO) vs. PMC78 150 μM | \*\*\*\* | <0,0001 |
| Neg. Control (w/ 0.3% DMSO) vs. PMC78 250 μM | \*\*\*\* | <0,0001 |
| Neg. Control (w/ 0.3% DMSO) vs. PMC79 50 μM | ns | 0,1142 |
| Neg. Control (w/ 0.3% DMSO) vs. PMC79 150 μM | \*\* | 0,0031 |
| Neg. Control (w/ 0.3% DMSO) vs. PMC79 250 μM | \*\*\*\* | <0,0001 |
| Neg. Control (w/ 0.3% DMSO) vs. Adagrasib 100 μM | ns | 0,5859 |
| Time-point 8h | | |
| Neg. Control (w/ 0.3% DMSO) vs. PMC78 50 μM | ns | 0,8915 |
| Neg. Control (w/ 0.3% DMSO) vs. PMC78 150 μM | ns | 0,1874 |
| Neg. Control (w/ 0.3% DMSO) vs. PMC78 250 μM | \* | 0,0128 |
| Neg. Control (w/ 0.3% DMSO) vs. PMC79 50 μM | ns | 0,9994 |
| Neg. Control (w/ 0.3% DMSO) vs. PMC79 150 μM | ns | 0,3172 |
| Neg. Control (w/ 0.3% DMSO) vs. PMC79 250 μM | \*\*\*\* | <0,0001 |
| Neg. Control (w/ 0.3% DMSO) vs. Adagrasib 100 μM | ns | 0,1529 |
| Time-point 24h | | |
| Neg. Control (w/ 0.3% DMSO) vs. PMC78 50 μM | \*\* | 0,0037 |
| Neg. Control (w/ 0.3% DMSO) vs. PMC78 150 μM | ns | 0,1436 |
| Neg. Control (w/ 0.3% DMSO) vs. PMC78 250 μM | ns | 0,1834 |
| Neg. Control (w/ 0.3% DMSO) vs. PMC79 50 μM | ns | 0,2801 |
| Neg. Control (w/ 0.3% DMSO) vs. PMC79 150 μM | ns | 0,9880 |
| Neg. Control (w/ 0.3% DMSO) vs. PMC79 250 μM | \*\*\*\* | <0,0001 |
| Neg. Control (w/ 0.3% DMSO) vs. Adagrasib 100 μM | ns | 0,0579 |

### Slide 11
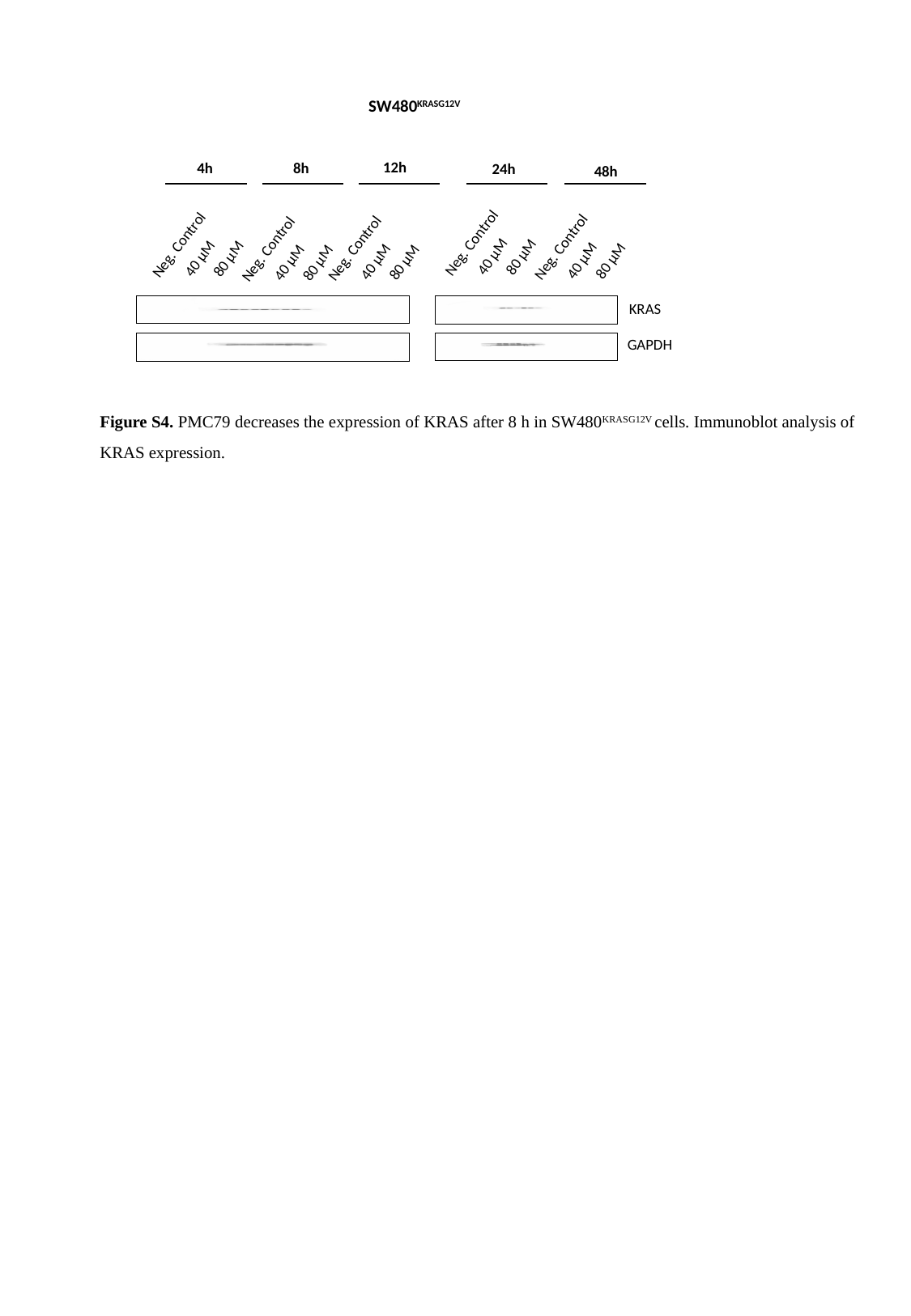

SW480^KRASG12V^
12h
8h
4h
24h
48h
Neg. Control
Neg. Control
Neg. Control
Neg. Control
Neg. Control
40 μM
80 μM
40 μM
80 μM
40 μM
80 μM
40 μM
80 μM
40 μM
80 μM
KRAS
GAPDH
Figure S4. PMC79 decreases the expression of KRAS after 8 h in SW480^KRASG12V^ cells. Immunoblot analysis of KRAS expression.

### Slide 12
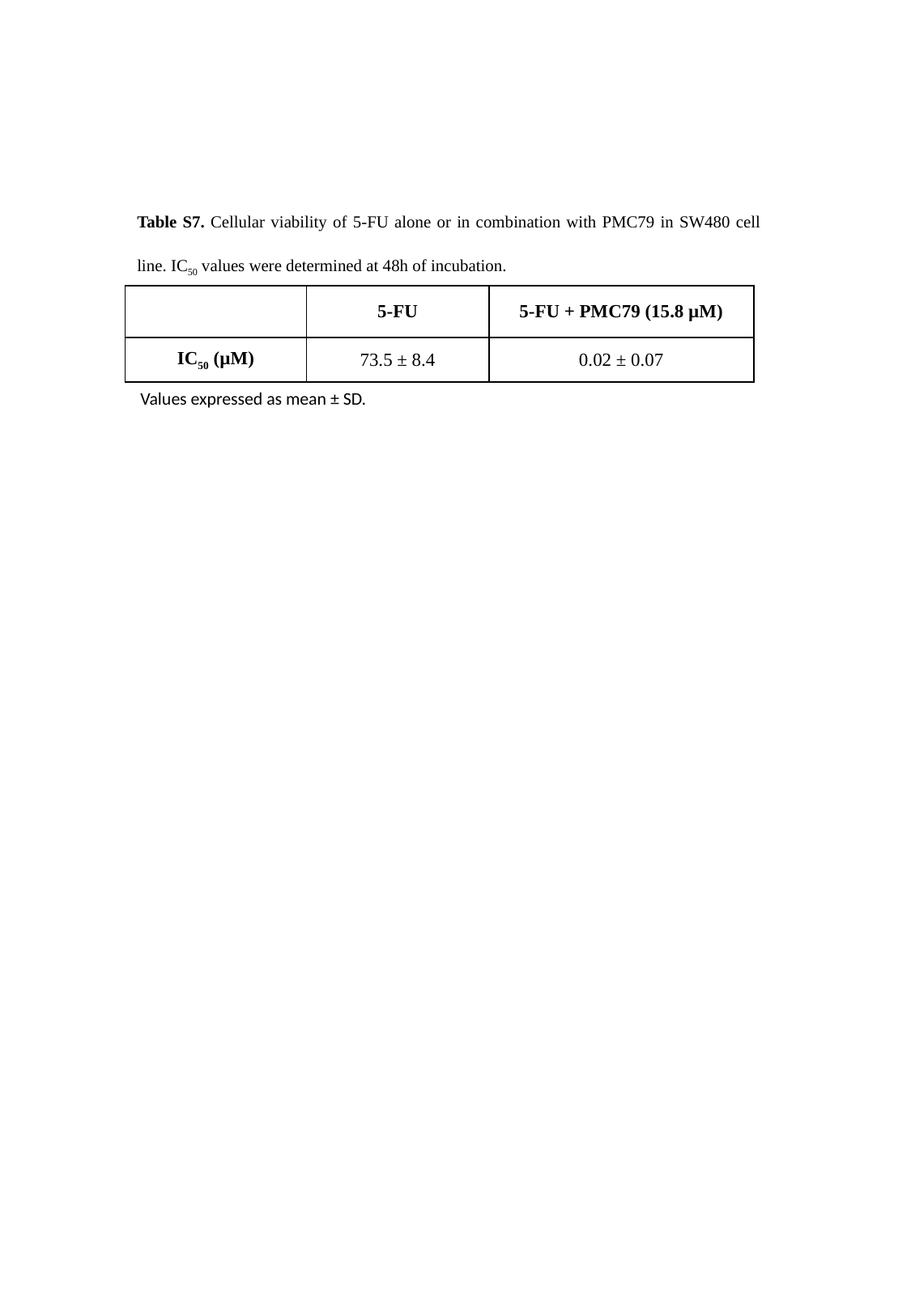

Table S7. Cellular viability of 5-FU alone or in combination with PMC79 in SW480 cell line. IC_50_ values were determined at 48h of incubation.
| | 5-FU | 5-FU + PMC79 (15.8 µM) |
| --- | --- | --- |
| IC_50_ (µM) | 73.5 ± 8.4 | 0.02 ± 0.07 |
Values expressed as mean ± SD.

### Slide 13
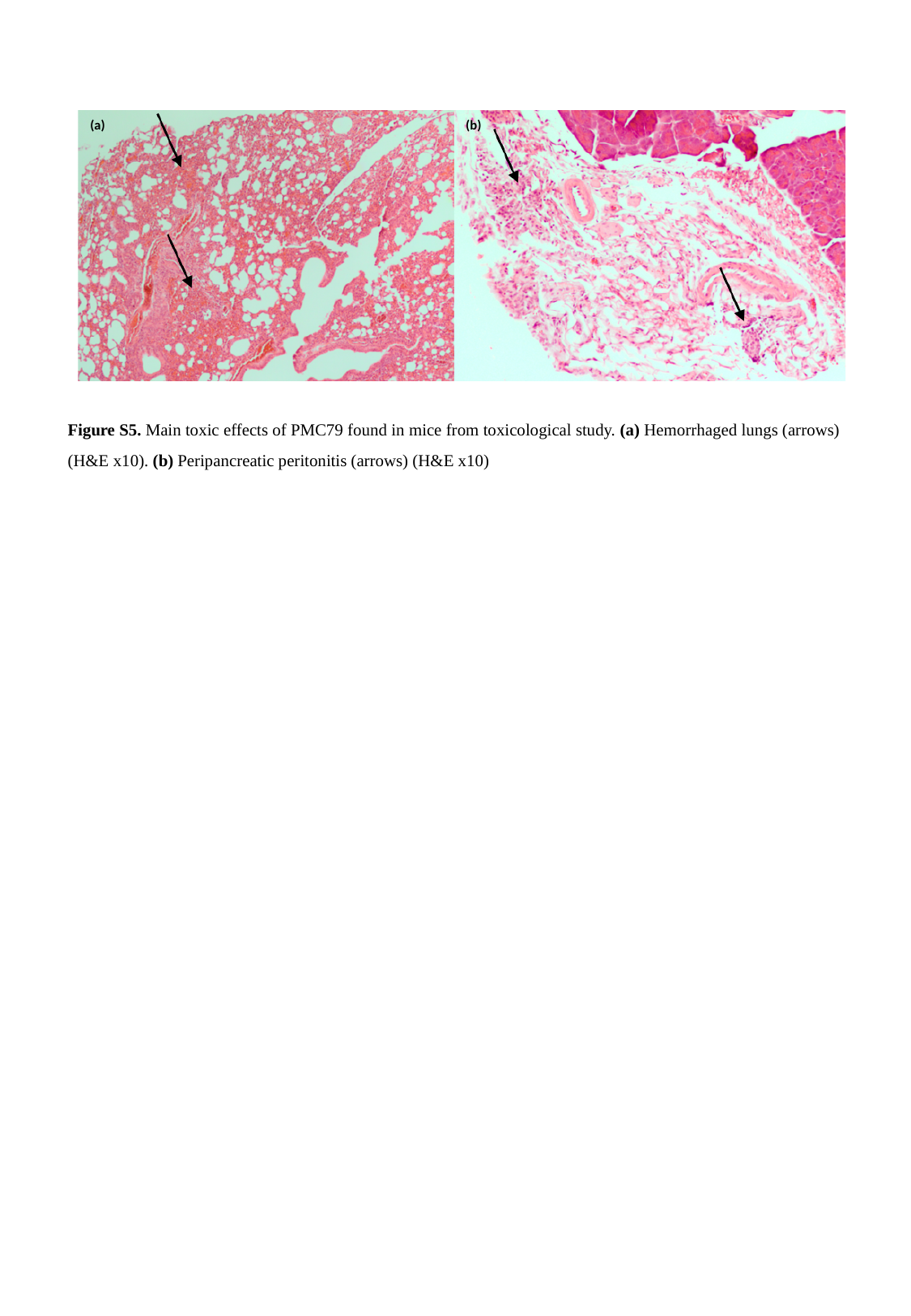

(b)
(a)
Figure S5. Main toxic effects of PMC79 found in mice from toxicological study. (a) Hemorrhaged lungs (arrows) (H&E x10). (b) Peripancreatic peritonitis (arrows) (H&E x10)

### Slide 14
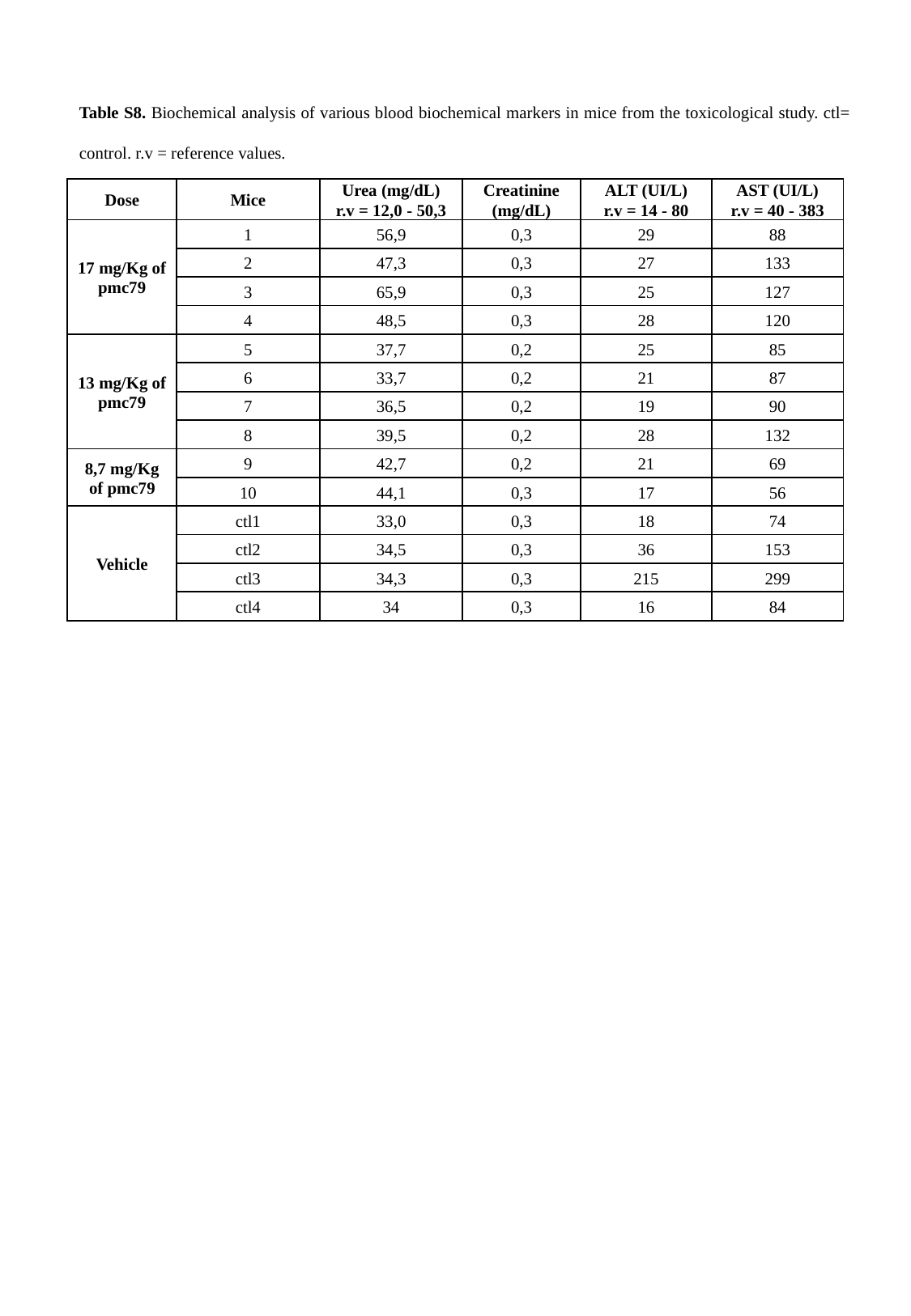

Table S8. Biochemical analysis of various blood biochemical markers in mice from the toxicological study. ctl= control. r.v = reference values.
| Dose | Mice | Urea (mg/dL) r.v = 12,0 - 50,3 | Creatinine (mg/dL) | ALT (UI/L) r.v = 14 - 80 | AST (UI/L) r.v = 40 - 383 |
| --- | --- | --- | --- | --- | --- |
| 17 mg/Kg of pmc79 | 1 | 56,9 | 0,3 | 29 | 88 |
| | 2 | 47,3 | 0,3 | 27 | 133 |
| | 3 | 65,9 | 0,3 | 25 | 127 |
| | 4 | 48,5 | 0,3 | 28 | 120 |
| 13 mg/Kg of pmc79 | 5 | 37,7 | 0,2 | 25 | 85 |
| | 6 | 33,7 | 0,2 | 21 | 87 |
| | 7 | 36,5 | 0,2 | 19 | 90 |
| | 8 | 39,5 | 0,2 | 28 | 132 |
| 8,7 mg/Kg of pmc79 | 9 | 42,7 | 0,2 | 21 | 69 |
| | 10 | 44,1 | 0,3 | 17 | 56 |
| Vehicle | ctl1 | 33,0 | 0,3 | 18 | 74 |
| | ctl2 | 34,5 | 0,3 | 36 | 153 |
| | ctl3 | 34,3 | 0,3 | 215 | 299 |
| | ctl4 | 34 | 0,3 | 16 | 84 |
